# Nucleocapsid condensation drives Ebola viral factory maturation and dispersion

**DOI:** 10.1101/2023.11.06.565679

**Authors:** Melina Vallbracht, Bianca S. Bodmer, Konstantin Fischer, Jana Makroczyova, Sophie L. Winter, Lisa Wendt, Moritz Wachsmuth-Melm, Thomas Hoenen, Petr Chlanda

## Abstract

Replication and genome encapsidation of many negative-sense RNA viruses take place in virus-induced membrane-less organelles termed viral factories (VFs). While liquid properties of VFs are believed to control the transition from genome replication to encapsidation, the nucleocapsid assembly, VF maturation and interactions with the cellular environment remain elusive. Here we apply *in situ* cryo-correlative light and electron tomography to follow nucleocapsid assembly and changes in VF morphology and their liquid properties during Ebola virus infection. We show that Ebola viral nucleocapsids transition from loosely packed helical assemblies in early VFs to condensed cylinders that arrange into highly organized parallel bundles later in infection. Early VFs associate with intermediate filaments and are devoid of other host material, but become progressively accessible to cellular components. Our data suggest that this process is coupled to VF solidification and dispersion, and that changes in liquid properties of VFs promote nucleocapsid transport to budding sites.

**Highlights:**

- Cryo-ET reveals the molecular architecture of Ebola virus replication compartments
- Loosely coiled nucleocapsids transition to condensed cylinders forming bundles
- Nucleocapsid condensation drives dispersion of viral factories promoting viral egress
- Intermediate filaments associate with and are critical for virus factory formation

## Introduction

Liquid-liquid phase separation has emerged as a fundamental organizing principle of cell compartmentalization, which can also be exploited by viruses to segregate essential viral replication and assembly processes from cellular activities [1, 2]. Non-segmented negative strand RNA viruses (NNSVs), such as Ebola virus (EBOV), Nipah virus, rabies virus, measles virus and respiratory syncytial virus, represent some of the most important human viruses. A hallmark of infections by these viruses is the formation of cytoplasmic biomolecular condensates that serve as viral replication factories (VFs). These structures provide a protective membrane-less microenvironment for efficient viral transcription and replication shielded from recognition by the host immune system [3–5] and in some cases enable persistent viral infection that can be reactivated by stress factors [6].

EBOV replication and transcription are confined to VFs also termed inclusion bodies [7], where the viral RNA and the replication machinery components, nucleoprotein (NP), polymerase L, transcriptional activator viral protein 30 (VP30), and polymerase cofactor VP35, concentrate. Recent work showed that EBOV VFs are *bona fide* liquid organelles [8–10] harnessing viral polymerase activity [7] and maintaining integrity through NP-NP interactions even in the absence of the viral RNA genome [8]. EBOV NP is considered the main driving force for liquid organelle formation and facilitates the recruitment of VP35 [11, 12], VP30 [13], VP24 [14] and L [15, 16] into the VFs. In analogy to the viral phosphoprotein P specific to the majority of NNSVs, EBOV VP35 is proposed to tether L to NP that form a replication-competent ribonucleoprotein complex. At later stages of infection, the matrix protein VP40 is also localized to VFs [17].

Besides viral RNA transcription and replication, conventional electron microscopy (EM) studies on sections of chemically fixed cells suggest that EBOV VFs also orchestrate encapsidation of the 19 kb long genome into a nucleocapsid (NC) [18]. However, all previous structural studies were performed on purified virions or virus-like particles and the process of NC assembly directly inside the VFs remains poorly understood. *In vitro* studies have demonstrated that binding of EBOV NP to RNA can assemble a loosely coiled NC-like structure [19], which *in vivo* presumably provides access to the polymerase L in association with VP35 to carry out replication and transcription [15, 20]. Previous EM and *in vitro* studies revealed that NCs undergo major structural rearrangements upon expression of VP40, VP35 and the NC-associated protein VP24 [18, 19]. This involves the transition of the loosely coiled helical assemblies into rigid thick-walled cylindrical NCs resembling those present inside EBOV virions whose structure was recently determined by cryo-ET [19, 21]. VP24 and VP35 have been mapped into the appendages at the outer surface of the NC and are proposed to facilitate condensation of loosely coiled helical NP oligomers [21, 22]. Condensed NCs are thought to undergo actin-dependent trafficking towards the plasma membrane [23, 24], where they are incorporated into budding virions formed by the VP40 matrix layer and the viral glycoprotein GP [18, 25].

While much of the attention has been dedicated to the function of VFs, there remains a significant gap in our understanding of their development and physical properties during the course of viral infection. Previous studies have demonstrated that viral biomolecular condensates can coordinate virus capsid assembly [4, 5, 26] and recent models suggest that phase separation accelerates and prevents kinetic trapping of capsid assembly [27]. However, the connection between RNA replication, NC assembly and physical properties of the VFs is not understood and has not yet been investigated on a molecular level. Identifying ultrastructural changes of VFs and how they relate to their liquid properties during infection will inform on the spatio-temporal coordination of viral replication and maturation of NCs. Moreover, the EBOV NC structure and the sequence of events leading to its assembly have not been studied directly in infected cells.

Here we aim to address these knowledge gaps by combining cellular cryo-electron tomography (ET) and *in situ* fluorescence recovery after photobleaching (FRAP) of authentic EBOV VFs. By employing *in situ* cryo-correlative fluorescence microscopy, cryo-ET and subtomogram averaging, we elucidate the structural rearrangements of the EBOV replication compartment and the NC assembly cascade at high resolution in cells infected with authentic EBOV. We show that NC condensation controls the fluidity of the VFs. The NC assembly and corresponding decrease in VF fluidity eventually leads to VF dispersion in the cytosol and liberation of the NC, thereby rendering it available for interaction with the cytoskeleton and permitting its trafficking towards viral budding sides.

## Results

### EBOV viral factories undergo morphological maturation concomitant with nucleocapsid condensation and bundle formation

To elucidate the morphological changes of EBOV VFs over time we infected Huh7 cells with wildtype EBOV, chemically fixed them at different time points post infection (p.i.) and visualized the VFs by confocal immunofluorescence microscopy using an antibody against EBOV NP, the main driving force for VF formation (Fig. 1A) [8]. VFs of each time point were segmented (Fig. 1A, middle panel) and analyzed with respect to their shape (sphericity) (Fig. 1A, B), surface area and volume (Fig. 1C-G). Early in infection (14 h p.i.) VFs revealed a spherical shape (sphericity of 0.85) with an average volume of the largest VF in each cell of 47 µm^3^. As infection progresses, VFs grow to larger condensates with an average maximum volume of 240 µm^3^ and 502 µm^3^ at 18 h p.i. and 22 h p.i., respectively. Concomitant with the increase in VF size we observed a reduction in sphericity dropping from 0.85 at 14 h p.i. to 0.77 and 0.78 at 18 h and 22 h p.i., respectively, suggesting a change in the liquid properties of the VFs. The VF morphology changed from spherically well-separated droplet-like structures to rugged structures of variable shapes dispersed across the cytosol and an increasing number of NP punctate structures was observed (Fig 1A). Overall, EBOV VFs become less equally sized during the course of infection, as indicated by an increasing Gini coefficient (Fig. 1D), highlighting changes in their dynamic character, shape and size.

**Fig. 1:**
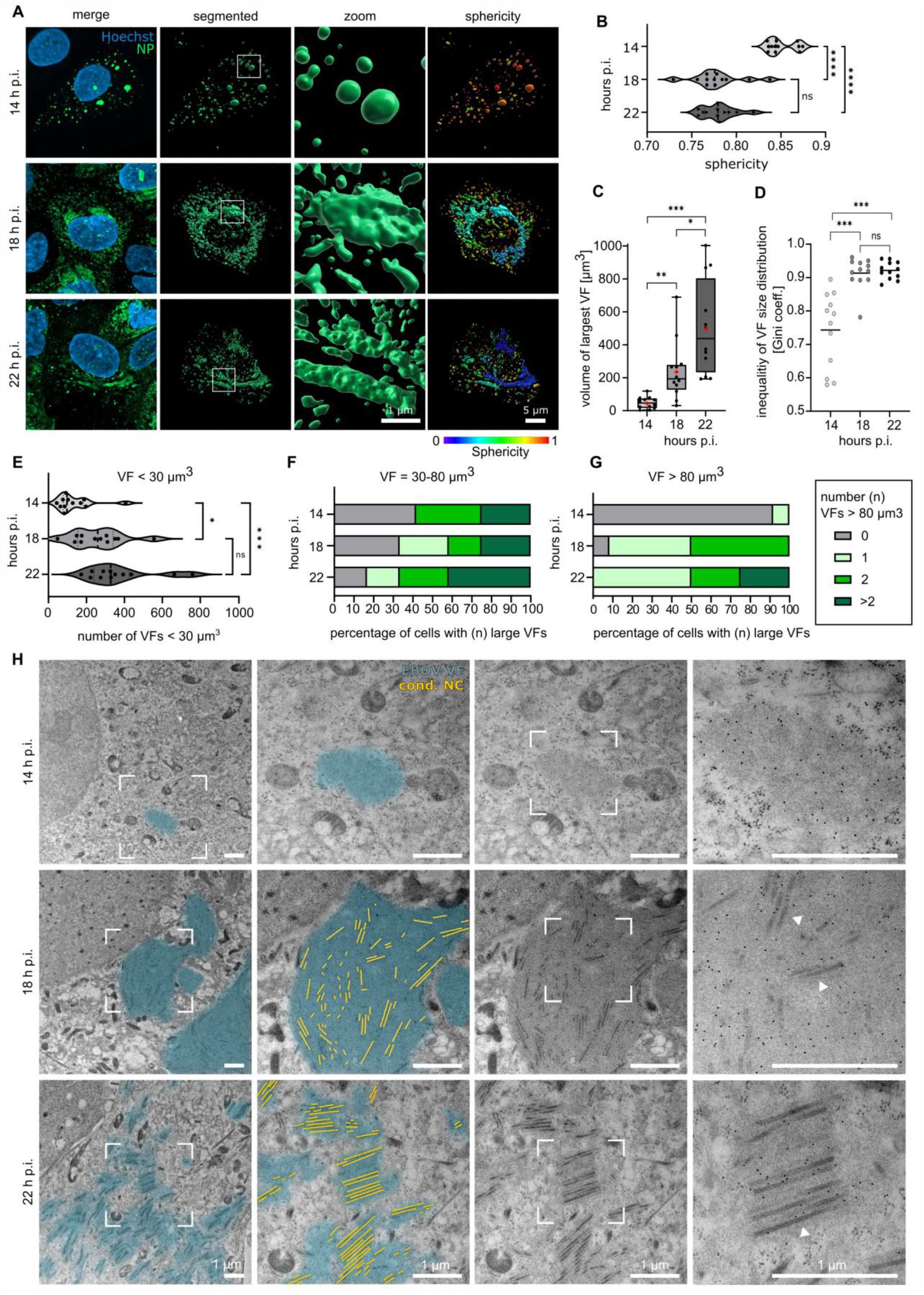
Morphological changes of EBOV viral factories during the course of an infection. **(A)** Confocal microscopy analysis of wildtype EBOV infected cells fixed at 14, 18 and 22 h p.i. Cells were immuno-stained against the EBOV NP forming the viral factories (VFs) (green). Nuclei were stained with Hoechst (blue). Maximum-intensity projections of representative cells for all time points (first panel) and three-dimensional segmentation of VFs and respective enlarged views (middle panels) are shown. Individual objects are color-coded according to their sphericity (right panel). **(B-G)** Quantification of VFs. At the indicated time points p.i. the VF sphericity **(B)**, volume of the largest VF **(C)**, inequality of VF size distribution **(D)**, the number of small VFs (volume < 30 µm^3^) **(E)**, intermediate VFs (volume between 30-80 µm^3^) **(F)**, and large VFs (volume > 80 µm^3^) **(G)** per cell was quantified. n = 4 cells per time point. Red plus in **(C)** marks the mean. Statistical significance was evaluated using a two tailed Welch’s t-test. ns; not significant, ∗: p < 0.1, ∗∗: p < 0.01, ∗∗∗: p < 0.001, and ∗∗∗∗: p < 0.0001. **(H)** Thin-section transmission electron microscopy analysis of EBOV VFs (blue) at indicated time points p.i. identified by immuno-labelling of EBOV NP. Condensed nucleocapsids (cond. NCs, yellow) are first visible at 18 h p.i. and form large bundles at 22 h p.i. concomitant with VF dispersion. Section thickness: 100 nm.

We next employed thin-section transmission electron microscopy (thin-section TEM) to examine morphological changes of EBOV VFs at the ultrastructural level (Fig. 1H). To this end, Huh7 cells were infected with wildtype EBOV, chemically fixed at 14 h, 18 h and 22 h p.i. and subjected to high pressure freezing and freeze substitution. The area of VFs was determined by immuno-EM on 100 nm-thick resin sections using an EBOV NP specific antibody. At early time points (14 h p.i.) EBOV VFs were detectable as small (< 5 µm in diameter), round electron-dense regions in cytoplasmic areas. At 18 h p.i. the area of the NP-positive, electron-dense regions increased and contained electron-dense filamentous structures exhibiting a central channel likely corresponding to assembled, condensed EBOV NCs (Fig. 1H, Fig S1A). Condensed NCs were distributed throughout the whole VF (Fig. 1H, 18 h p.i.), indicating that NC assembly occurs across all regions within the VF, without a specific VF subregion being favored. In addition to many individual NCs, we observed some bundles of 2-5 NCs arranged with parallel alignment at 18 h p.i (Fig. S2). NC bundling became more prominent at late times post-infection (22 h p.i.), when larger bundles (> 5 NCs per bundle) with up to 21 NCs were detected (Fig. S2). Concomitant with the formation of NCs into larger bundles at 22 h p.i., we observed a decrease of NP positive electron-dense areas and dispersion of the VFs into smaller areas. This data suggests that the increasing number of small NP positive spots observed in immunofluorescence analysis at 22 h p.i. (Fig. 1A, E) correspond to VFs containing bundles of parallel arranged NCs resulting from VF dispersion. Thus, as the infection progresses, VFs disperse and form individual and partially interconnected compartments each containing at least one bundle of condensed NCs (Fig. 1H, lower panel). Large bundles of NCs were also observed near the plasma membrane where budding takes place (Fig. S1B, C, region a-b). However, only individual NCs were found horizontally aligned with the plasma membrane, showing that separation of the bundles likely precedes virion budding (Fig. S1C, region a,c). In summary, our data demonstrate that NC condensation takes place in VFs as early as 18 h p.i., and uncovers that VFs undergo dispersion at late time points in infection likely as a result of NC condensation and bundle formation.

### NC maturation involves condensation of loosely coiled helical assemblies

To investigate the structural determinants of VF maturation at higher resolution, we applied cellular cryo-ET and took advantage of an rgEBOV-VP30-GFP reporter virus, which forms VFs containing VP30-GFP without compromising VF function [8, 28]. This allowed us to localize VFs by cryo-confocal microscopy directly on cryo-focused ion beam (cryo-FIB) milled lamellae and acquire cryo-electron tomograms in a correlative fashion (Fig. 2). Cells analyzed by cryo-ET at 14 h p.i. showed that VFs are exclusively composed of loosely coiled left-handed helical structures (Fig. 2A-F) resembling those that assemble *in vitro* upon expression of NP and represent the precursor to the fully assembled EBOV NC [29, 30]. The results could be confirmed by cryo-ET analysis of cells infected with wildtype EBOV fixed at 14 h p.i. (Fig. 3A-E, Movie S1). The loosely coiled NCs had an average diameter of approximately 34 nm, a pitch of 24 nm (Fig. 3E) and were often curved, indicating that they are highly flexible. In accordance with the EM data (Fig. 1H), cryo-ET analysis of VFs at 18 h and 22 h p.i. revealed, in addition to loosely coiled NCs, straight cylindrical structures with an outer diameter of 50 nm and an inner diameter of 20 nm (Fig. 2G-L, Fig. 3F-N, Movie S2). Overall, the data shows that early after infection, VFs are mainly composed of loosely coiled NCs which upon interaction with other viral components condense to form a thick-walled rigid cylinder in the cytoplasmic VFs at later stages of infection.

**Fig. 2.**
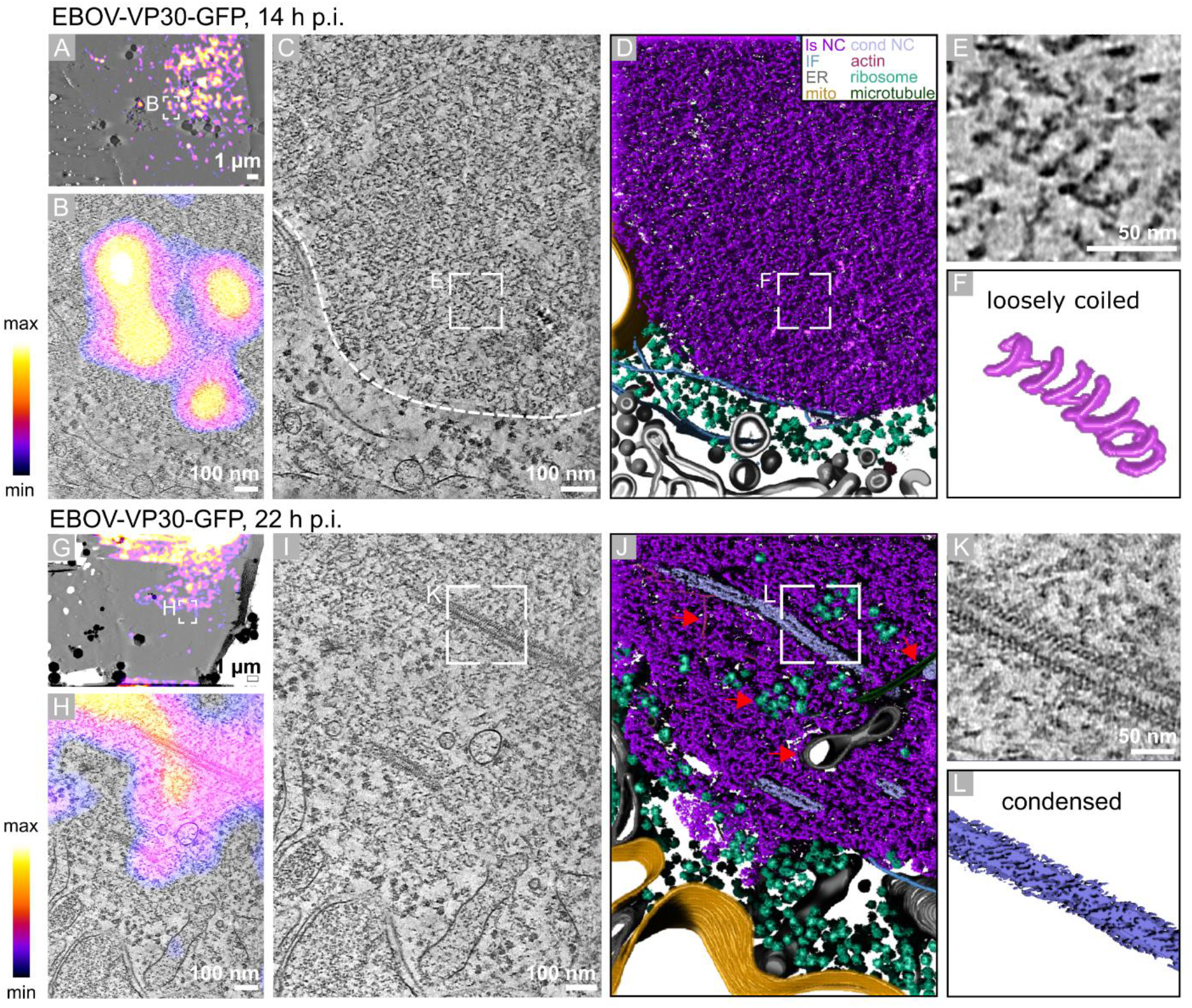
On lamella cryo-correlative light and electron tomography identifies viral factories in EBOV-VP30-GFP infected cells. **(A, G)** On lamella cryo-correlative light and electron tomography analysis of cells infected with rgEBOV-VP30-GFP and fixed at 14 h **(A)** and 22 h p.i. **(G)**. Overlay of a cryo-TEM image of lamella (grey) with cryo-Airyscan fluorescence signal of VP30-GFP (fire lookup table). Areas where tomograms were acquired are indicated. **(B, H)** Tomographic slice (grey) of rgEBOV-VP30-GFP infected cells at 14 h p.i. (B) and 22 h p.i. (H) overlaid with cryo-Airyscan fluorescence signal of VP30-GFP (fire lookup table). **(C, I)** Tomographic slice showing the same area as in **(B, H)**. Dotted line in **(C)** indicates the border of the viral factory. **(D, J)** Segmentation of the tomograms shown in **(C, I)**. Color legend indicates the observed viral and cellular components; loosely coiled nucleocapsid (ls NC) (purple), condensed nucleocapsid (cond NC) (violet), intermediate filament (IF) (blue), actin (red), endoplasmic reticulum (ER) (grey), ribosome (green), mitochondrium (mito) (yellow), microtubule (dark green). Cellular structures invading the space of the VFs are highlighted by red arrows **(J)**. **(E, F, K, L)** Close-up views of a loosely coiled **(E)** and a condensed nucleocapsid **(K)** and corresponding segmentations **(F, L)**.

**Fig. 3.**
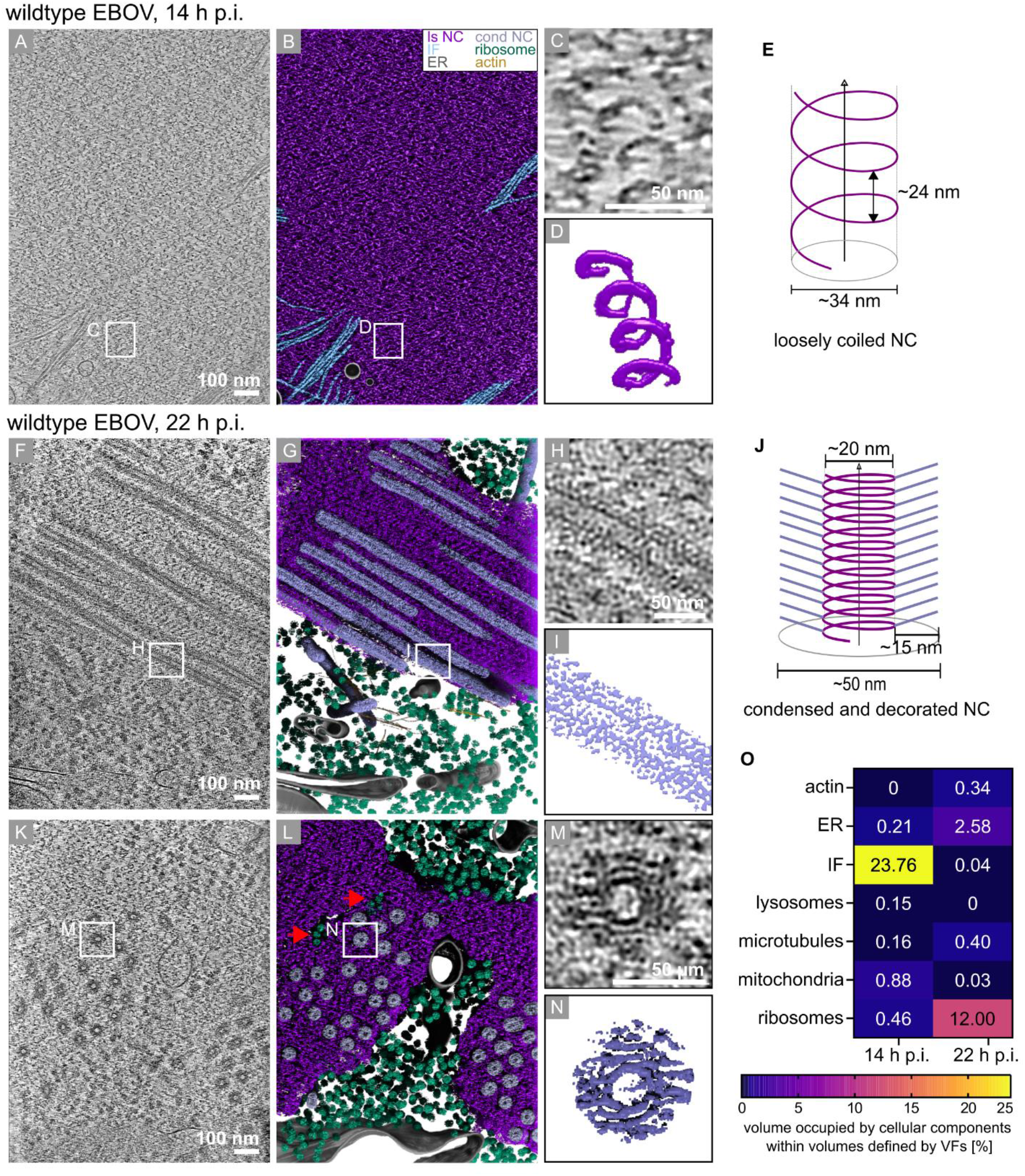
Molecular architecture of wildtype EBOV viral factories and interactions with cellular components. **(A**) *In situ* cryo-ET analysis of wildype EBOV infected cells at 14 h p.i. Tomographic slice showing that viral factories at 14 h p.i. are composed of loosely coiled left-handed helical nucleocapsids (NCs). **(B)** Segmentation of the tomogram shown in **(A)**. Color scheme indicates the observed viral and cellular components; loosely coiled nucleocapsid (ls NC) (purple), intermediate filament (IF) (blue), endoplasmic reticulum (ER) (grey), ribosome (green), actin (yellow). **(C, D)** Close-up view of a loosely coiled NC **(C)** and corresponding segmentation **(D)**. **(E)** Model of a loosely coiled NC. Average diameter and pitch of the helical assembly are indicated. **(F, K)** Slice through a tomogram of wildtype EBOV infected cells at 22 h p.i. showing that viral factories contain parallel bundles of condensed NCs. Side views **(F)** and top views **(K)** of NCs are shown. **(J)** Model of a condensed NC. Average outer and inner diameter and length of the decoration are indicated. **(G, L)** Segmentations of the tomograms shown in **(F, L).** Color scheme indicating the observed viral and cellular components according to **(B)**. Ribosomes invading the space of the VFs are highlighted by red arrows **(L)**. **(H, I, M, N)** Close-up views of a condensed NC from side **(H)** and top view **(M)** and corresponding segmentations **(M, N)**. **(O)** Volume quantification of cellular components found within the volume defined by VFs at 14 h p.i. and 22 h p.i. VFs, i.e. volume occupied by both loosely coiled and condensed NCs, were segmented and the respective volume was set as 100%. Cellular structures observed within these volumes were segmented. The heat map indicates how much volume of the VF is occupied by the indicated cellular components at the indicated time points p.i. (n = 4 cells for each time point).

### NCs of defined lengths organize into bundles arranged in a hexagonal lattice

To obtain structural details of the intracellular condensed EBOV NC, we applied subtomogram averaging using 4,426 particles extracted from 8 tomograms acquired on lamellae of wildtype EBOV infected cells at 22 h p.i. (Fig. S3A). This yielded a map at 21 Å resolution (0.143 cut-off) revealing a left-handed helical tube with a hollow smooth-walled interior with a diameter of 20 nm (Fig. 4A, B, S3B). The outer wall of the helical assembly was decorated with appendages and exhibited an outer diameter of 50 nm, a pitch of 7.5 nm and 12 units per turn (Fig. 4A, B). This structure agreed well with the structure of the condensed EBOV NC in isolated virions previously determined by subtomogram averaging [21]. Hence, it represents a mature condensed NC which presumably encapsidates the 19 kb EBOV RNA genome of negative polarity. To investigate whether the condensed NCs found inside the VFs could accommodate the EBOV genome, we measured the length of complete NCs horizontally oriented within the lamellae. While the NC length was variable at 18 h p.i. ranging from 390-930 nm, NCs measured at 22 h p.i. had a consistent length of 918 nm ± 171 nm (mean and standard deviation) (Fig. 4C). Based on previously published data showing that EBOV packages 6 RNA nucleotides per copy of NP into a helical assembly with a pitch of ∼7.5 [31], the calculated length of the NC accommodating the 18959 nucleotide genome of EBOV is 911 nm. Thus, the average length of 918 nm measured for NCs at 22 h p.i. corresponds to the calculated length of the NC encapsidating one EBOV genome.

**Fig. 4.**
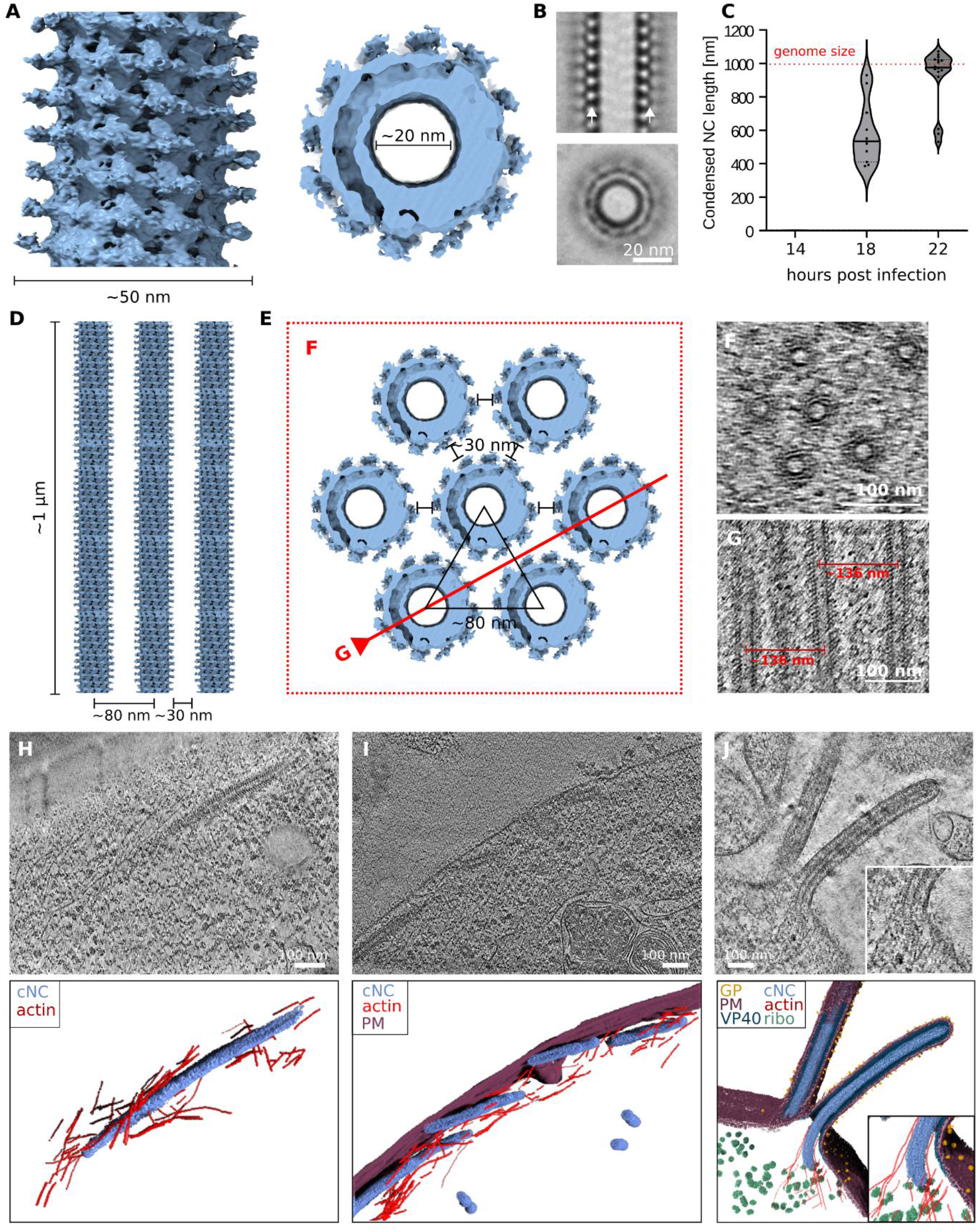
Structure, bundling and trafficking of the Ebola virus nucleocapsid in infected cells. **(A)** *In situ* structure of the condensed EBOV nucleocapsid (NC) from side (left panel) and top view (right panel) determined by subtomogram averaging of tomograms from wildtype EBOV infected cells at 22 h p.i. Outer and inner diameter of the NC are indicated. **(B)** Orthoslices taken through the NC structure determined by subtomogram averaging. White arrow heads indicate the decoration. **(C)** Length measurements of condensed NCs in cryo-lamellae prepared from wildtype EBOV infected cells at 14 h p.i. (not detected), 18 h p.i. (581 nm ± 178 nm) and 22 h p.i. (919 nm ± 171 nm). n = 11 NCs for 18 h p.i.; n = 16 NCs for 22 h p.i. Red dotted line indicates the length of the 19 kb EBOV genome considering that one NP binds to six RNA nucleotides. **(D-G)** Model of EBOV NC parallel bundling at 22 h p.i. Mature condensed NCs with a length of ∼1 µm organize into bundles arranged in a hexagonal lattice. The minimal average distance between the NCs measured from center-to-center and between the outer protrusions is indicated. Model is not drawn to scale **(D, E)**. Red dotted box and line mark the positions of the cross sections shown in **(F, G)**. Slices through representative tomograms of wildtype EBOV infected cells at 22 h p.i. showing NC bundles from top **(F)** and side view **(G)**. Center-to-center distance between two NCs is indicated **(G)**. **(H-J)** *In situ* cryo-ET of wildype EBOV infected cells at 22 h p.i. Tomographic slices (upper panel) and corresponding 3D segmentations (lower panel) are shown. Color code: condensed nucleocapsid (cNC) (blue), actin filament (red), plasma membrane (PM) (berry), VP40 (dark blue), glycoprotein (GP) (yellow), ribosome (ribo) (green). Individual transport-competent NC in the cell periphery associated with actin filaments **(H)**. NCs aligning at the PM for budding **(I)**. Incorporation of a NC into a budding virion **(J)**.

Cryo-ET allowed us to provide insights into the spatial organization of NCs within EBOV VFs. Tomograms of lamellae capturing both transverse and longitudinal cross-sections of condensed NCs revealed NC parallel bundling with quasi-hexagonal packing (Fig. 4D-G, Movie S2). The nearest neighbor NC-NC distance forming a parallel bundle was 30 nm (Fig. 4D, E).

### VF dispersion allows NC separation and association with actin for transport to viral budding sites

In contrast to VFs at 14 h p.i., we observed that VFs at 22 h p.i. lost their overall integrity and that the space occupied by VFs also contained ribosomes, occasionally small vesicles and increasingly also cytoskeletal components such as actin and microtubules (Fig. 2I, J (arrow), 3K, L (arrow) and 3O quantification). In addition, *in situ* cryo-ET of infected cells at late time points p.i. revealed separated condensed NCs that associate with actin at the plasma membrane and condensed NCs in the process of incorporation into budding virions (Fig. 4H-J). Association of NCs with actin could be observed for the whole sequence of events taking place during budding, i.e. (i) the transport of NCs to the PM (Fig. 4H, Movie S3), (ii) lateral association of NCs with the plasma membrane (Fig. 4I), and (iii) NC envelopment at the plasma membrane (Fig. 4J). Overall, these data support a model in which VF dispersion leads to loss of VF integrity, which allows association of condensed NCs with actin filaments. This in turn facilitates NC transport to viral budding sites at the plasma membrane and subsequent incorporation into virions.

### EBOV VFs associate with cellular vimentin important for the viral life cycle

A main biological function of liquid organelles is the separation of an initially homogeneous mixture into components which accumulate preferably inside and outside the condensate, respectively. This can lead to a large increase of concentrations of certain components inside the condensates and exclusion of others, as shown before e.g. for centrosomes [32]. Using segmentation of the tomograms, we investigated the VFs for the enrichment of cellular components that can be identified by cryo-ET, such as ribosomes, cytoskeleton, membranous organelles and vesicles. Our data show that at 14 h p.i., VFs composed of loosely coiled NCs maintain high integrity and lack presence of most cellular structures (Fig. 2C, D, 3A, B and 3O quantification, Movie S1). Strikingly, the only visible cellular component that appeared to invade the VFs were filamentous structures with a diameter of ∼12 nm corresponding to vimentin intermediate filaments (Fig. 5A-E, 3O quantification, Movie S4). These observations were further supported by thin-section TEM analysis of EBOV infected cells (Fig. S4A-C) and confocal immunofluorescence microscopy using an anti-vimentin antibody (Fig. 5F, S4D). In mock-infected cells vimentin forms a filament network that is evenly distributed throughout the cytoplasm (Fig. S4D). Upon wildtype EBOV infection the localization of vimentin rearranges, and the intermediate filaments form compact bundles in the perinuclear region near the VFs without any notable changes in overall cell morphology (Fig. S4D, Fig. 5F). 3D segmentation of the confocal data revealed vimentin in close proximity to VFs, occasionally present in their interior (Fig. 5F, left panel). We next asked whether vimentin plays a role in VF biogenesis and integrity. Treatment of transfected cells with withaferin A (WFA), which selectively disrupts the vimentin cytoskeleton, prevented the formation of large VFs (Fig. 5F, G), indicating that vimentin plays a role in VF growth. Finally, to test whether the intact vimentin cytoskeleton supports the EBOV replication cycle, we evaluated RNA synthesis and protein expression of EBOV in the presence or absence of WFA used in concentrations below cytotoxic conditions using an EBOV reporter virus expressing firefly luciferase (rgEBOV-luc2) [33] (Fig. 5H, I). Analysis of luciferase activity showed that WFA significantly inhibits EBOV RNA synthesis and/or protein expression. Overall, these data show that VFs associate with vimentin and indicate that vimentin supports the growth of these structures.

**Fig. 5.**
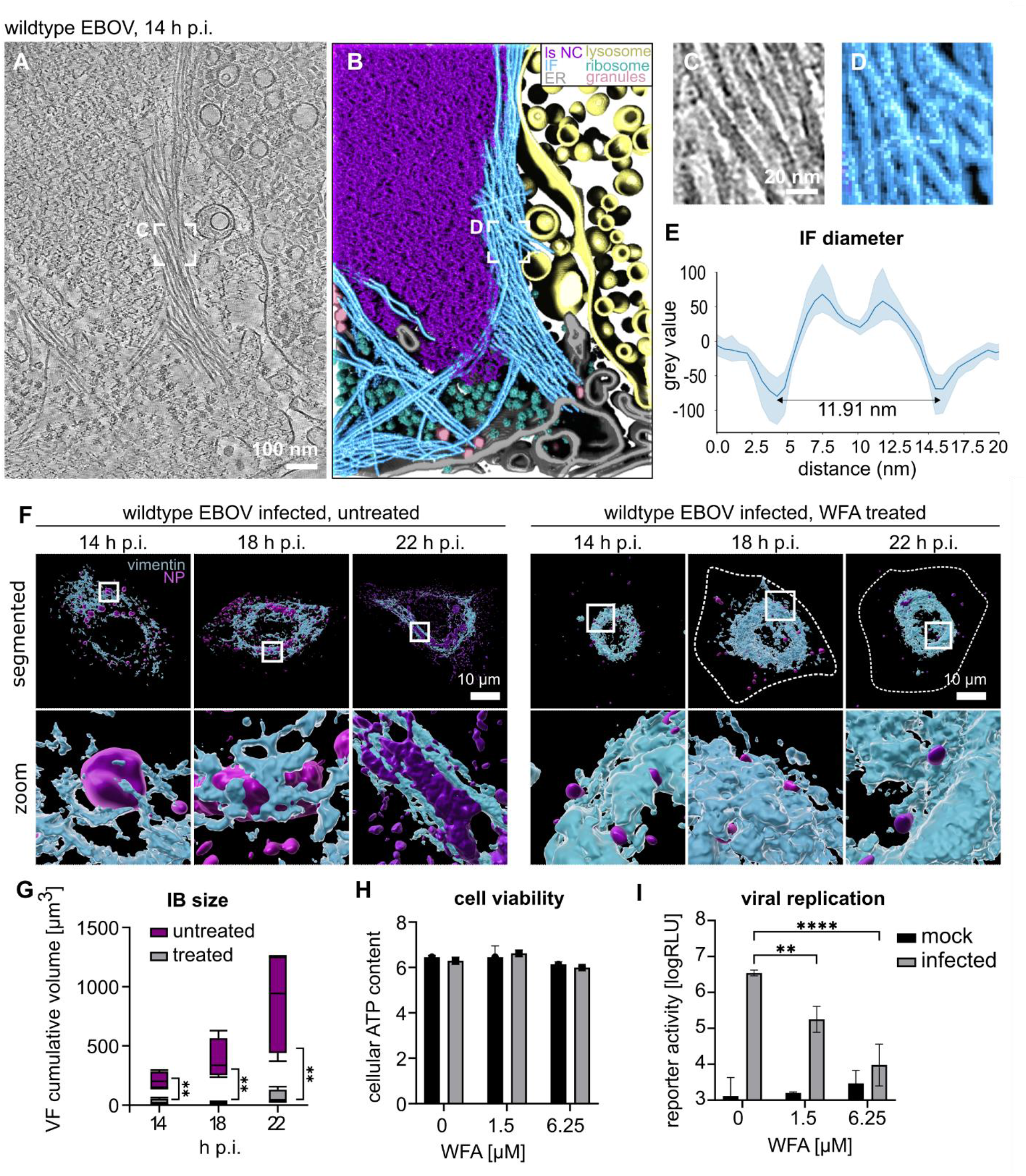
EBOV VFs associate with vimentin intermediate filaments important for the viral life cycle. **(A)** Slice through a representative tomogram of a cell infected with wildtype EBOV and fixed at 14 h p.i. **(B)** Segmentation of the tomogram shown in **(A)**. Color scheme indicates the observed viral and cellular components; loosely coiled nucleocapsid (ls NC) (purple), intermediate filament (IF) (blue), endoplasmic reticulum (ER) (grey), lysosome (yellow), ribosome (green), granules (pink). **(C, D)** Close-up views of IFs. **(E)** Average normalized density line profile of IF cross-sections. Error band shows the standard deviation (n = 20 from four tomograms). **(F)** Confocal microscopy analysis of wildtype EBOV infected cells fixed at 14, 18 and 22 h p.i. Cells were immuno-stained against the EBOV nucleoprotein forming the viral factories (VFs) (purple) and against vimentin intermediate filaments (light blue). Shown are three-dimensional segmentations of the VFs and vimentin (upper panel) and respective enlarged views (lower panels). Right panels show 3D segmentations of cells treated with withaferin A (WFA) at a concentration of 6.25 µM. Dotted line indicates plasma membrane. **(G)** Quantification of VF sizes at 14, 18 and 22 h p.i. with wildtype EBOV with and without treatment with WFA at the indicated concentrations. Statistical significance was evaluated using two tailed Welch’s t-test. ∗∗: p < 0.01. **(H)** Cell viability of cells infected with EBOV or left uninfected and treated with WFA or left untreated (mock DMSO control). Cell viability was determined by quantification of intracellular ATP levels. **(I)** Effect of WFA on EBOV replication. Cells were infected with rgEBOV-luc2 and treated with indicated concentrations of WFA or left untreated (DMSO control, mock). One day p.i. luciferase activity in relative light units (RLU) was measured. Means and standard deviations from three independent experiments are shown. Statistical significance was evaluated using two-way Anova, Dunnett’s multiple comparisons test. ∗∗: p < 0.01, and ∗∗∗∗: p < 0.0001.

### Nucleocapsid condensation modulates the fluidity of EBOV VFs

Our data showed that VFs undergo dramatic morphological changes in the course of EBOV infection and that this is concomitant with NC condensation, VF dispersion and NC interaction with actin. Based on these results we hypothesized that the liquid properties of VFs are altered at late time points of infection, marking the transition from genome replication to the assembly and egress phase of infection. To investigate the liquid properties of VFs we performed fluorescence recovery after photobleaching (FRAP) experiments of VFs formed at 14 h, 18 h and 22 h p.i. using rgEBOV-VP30-GFP infected cells (Fig. 6A). Consistent with our recently published results [8], the spherical VFs which dominate at 14 h p.i. showed a fast recovery of the VP30-GFP signal after photobleaching. However, both the recovery rate and efficiency of VFs formed at late times post-infection were decreased.

**Fig. 6.**
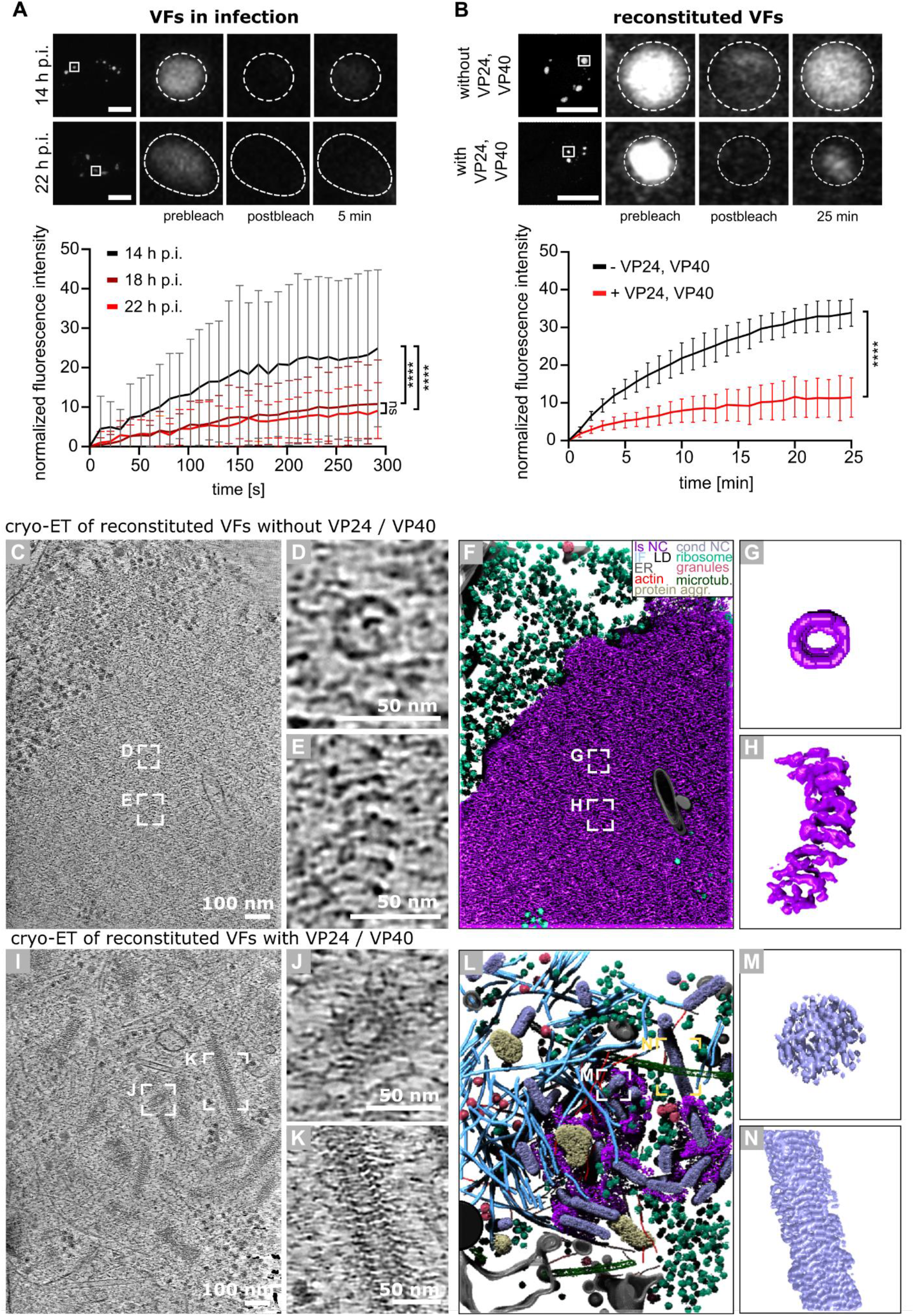
Nucleocapsid condensation modulates the fluidity of viral condensates. **(A)** Fluorescence images and quantification of fluorescence recovery after photobleaching (FRAP) analysis of EBOV VFs formed 14 h (black), 18 h (dark red) and 22 h (red) p.i. of cells with rgEBOV-VP30-GFP. Upper panel: Boxed areas indicate individual VFs chosen for photobleaching and zoom-in views shown on the right; dashed circles indicate bleached regions of VP30-GFP within VFs. Scale bars: 20 µm. Lower panel: Quantification shows the normalized mean fluorescence recovery curve and standard deviation (error bars). 14 h p.i. n = 27; 18 h p.i n = 38; 22 h p.i. n = 25. p values of differences analyzed by two-way ANOVA are shown. ns: not significant, ∗∗∗∗: p < 0.0001. **(B)** Fluorescence images and quantification of FRAP analysis of reconstituted EBOV VFs formed 28 h after transfection of cells with expression plasmids encoding EBOV NP, VP35, VP30-GFP and L (early-like VFs, black) or additionally VP24 and VP40 (late-like VFs, red). Representation as in **(A)**. Scale bars: 20 µm. Whole VFs were photobleached and fluorescence recovery was monitored. Quantification is similarly shown as in **(A).** Scale bars: 10 µm. n = 25 for both early-like and late like VFs. Statistical significance was evaluated using a two tailed Welch’s t-test. ∗∗∗∗: p < 0.0001. **(C-E)** *In situ* cryo-ET of reconstituted EBOV VFs formed 28 h post transfection of cells with expression plasmids encoding EBOV NP, VP35, VP30-GFP and L. Tomographic slice of a representative tomogram shows that VFs not containing VP24 and VP40 are exclusively composed of loosely coiled nucleocapsid-like structures **(C)**. Close-up view of a loosely coiled nucleocapsid from top **(D)** and side view **(E)**. **(F-H)** Segmentation of the tomogram shown in **(C)**. Color scheme indicates the observed viral and cellular components; loosely coiled nucleocapsid-like structures (ls NC) (purple), intermediate filament (IF) (blue), lipid droplet (LD) (black), endoplasmic reticulum (ER) (grey), actin (red), protein aggregate (protein aggr.) (light yellow), condensed NC (cond NC) (violet), ribosome (green), granules (pink), microtubules (microtub.) (dark green). Close-up view of the segmented loosely coiled nucleocapsid from top **(G)** and side view **(H)**. **(I-K)** *In situ* cryo-ET of reconstituted EBOV VFs formed 28 h post transfection of cells with expression plasmids encoding EBOV NP, VP35, VP30-GFP, L, VP40 and VP24. Tomographic slice of a representative tomogram shows that VFs containing VP24 and VP40 are composed of both loosely coiled and condensed NC-like structures **(I)**. Close-up view of a condensed NC from top **(J)** and side view **(K)**. **(L-N)** Segmentation of the tomogram **(L)**. Color scheme as in **(F)**. Close-up view of the segmented condensed nucleocapsid from top **(M)** and side view **(N)**.

To determine whether the decrease in VF fluidity is driven by NC condensation in the absence of the viral genome, we performed FRAP analysis of VFs that were reconstituted in cells using a transfection-based system (Fig. 6B). To mimic VFs at early stages of infection (early-like VFs) we transfected Huh7 cells with expression plasmids encoding EBOV NP, polymerase L, VP35 and VP30-GFP. Using *in situ* cryo-ET we could show that early-like VFs are indeed exclusively composed of loosely coiled NCs (Fig. 6C-H, Movie S5). In contrast, additional expression of VP24 and VP40, which are proposed to drive NC condensation [19, 21], led to the formation of VFs containing condensed NCs (Fig. 6I-N, Movie S6). In the absence of infection, NCs showed variable length ranging from 63-397 nm (Fig. S5) and did not form large parallel bundles (Fig. 6I-N). FRAP analysis of early-like and late-like VFs revealed that early-like VFs exhibit higher fluorescence recovery rates compared to late-like VFs (Fig. 6B). Taken together, these results demonstrate that VF maturation and morphology changes are accompanied by a reduction in condensate fluidity. Our data further suggest that the change in VF liquid properties is controlled by NC condensation, and we show that genomic RNA determines the length of condensed NCs, and that NC bundling is compromised in the absence of EBOV genomes.

## Discussion

Viral NCs are frequently considered as inert structural entities. Combining high-resolution cellular cryo-ET, cell biology and FRAP analysis, our study highlights the dynamics of viral condensates during maturation and plasticity of NC structures assembling inside them. Our study demonstrates that virus-induced biomolecular condensates undergo biophysical and structural changes during maturation that drive essential steps of virus infection such as replication, virus assembly and egress. Using EBOV as a model NNSV, we show that VFs are subjected to a maturation process manifested by a reduction in fluidity, loss of their sphericity and adoption of a higher physical order which is governed by structural changes of the viral NC.

Cellular cryo-ET of wildtype EBOV infected cells show that early after infection (14 h p.i.), VFs exclusively consist of NP oligomers assembling loosely coiled helical structures that are flexible and often intertwined. Our 3D segmentations revealed that these early VFs exhibit a high integrity largely excluding cellular structures such as ribosomes. This demonstrates that viral translation must take place outside of the condensates and requires export of transcripts to the VF periphery substantiating previous results on EBOV VFs [34]. Similarly, for rabies and mumps VFs it was shown that translation is not an integral part of their function [4, 6]. Based on the comparative cryo-EM and FRAP data of VFs formed at early and late stages of infection, we propose that the loosely coiled NCs represent an immature, RNA synthesis-competent state which contributes to and maintains the fluidity and integrity of the VF required for efficient RNA synthesis taking place inside the condensate [7]. While most cytoskeletal components are excluded from the early VFs, filamentous structures identified as vimentin intermediate filaments form a large network that associates with the VFs (Fig. 5A-F, S4, Movie S4). Inhibition of vimentin polymerization led to the formation of small-sized VFs and significant reduction of viral replication (Fig. 5I). This indicates the importance of vimentin for VF growth and is an initial proof-of-concept that targeting of cellular structures interacting with virus-induced biomolecular condensate may provide a viable antiviral strategy in drug development. Importantly, for Dengue virus and Zika virus vimentin was found to play a structural role in maintaining the integrity of the viral replication organelles [35, 36], raising the possibility that such a strategy might be viable against a broader spectrum of viruses.

As the infection progresses, initially loosely coiled NCs undergo a remarkable condensation by 40% in diameter (34 nm to 20 nm) and by 70% in helical pitch (from ∼24 to ∼7 nm) (Fig. 3, 4). Using cryo-ET of intracellularly reconstituted VFs, we could attribute this NC condensation to VP24 and VP40, substantiating previous *in vitro* data showing that VP24 is a key factor in NP condensation [19, 37]. Condensation likely leads to steric locking of the RNA genome inside the rigid helical assembly and consequently, halting L-mediated RNA synthesis. In line with that, functional studies showed that expression of VP24 prevents L-mediated RNA synthesis [14, 38, 39]. We hypothesize that only viral genomic RNA is encapsidated into condensed NCs, and that RNA of positive polarity remains associated with loosely coiled NP oligomers. However, the mechanism of this selectivity remains to be addressed. Our data further shows that the NC size is dictated by the number of nucleotides in the EBOV RNA genome (Fig 4 C), consistent with a previous cryo-EM study on purified EBOV virions [19]. This is further supported by our cryo-ET data of intracellularly reconstituted condensates, showing that condensed NCs produced in the absence of viral genome are highly variable in length (Fig. S5). Since the length of condensed NCs found inside VFs (Fig. 4C) corresponds well to the calculated size of the EBOV NC encapsidating the full EBOV RNA genome, we conclude that encapsidation of the viral RNA genome into a condensed NC is completed within the VFs and that only mature condensed NCs are transported to the budding sites.

Our subtomogram average of condensed NCs found in EBOV-infected cells at late stages of infection reveals a tightly packed left-handed helical assembly with protruding densities, structurally resembling NCs found in purified virions and VLPs [21]. Furthermore, cellular cryo-ET allowed us to investigate the spatial distribution of condensed NCs within VFs. Notably, we revealed that NCs locally organize in small hexagonal arrays forming parallel bundles with equidistant spacing. Although we occasionally observed loosely coiled NCs interspacing the condensed NC, in the majority of cases the volume between condensed NCs is occupied by material lacking high-order organization. Recently, it was shown that L is not homogeneously distributed within the VF in cells transfected with an EBOV minigenome system lacking VP24 [9]. Since VP35 interacts with L [15], we hypothesize that L could be present in the interspaced volume and stays associated with NCs for incorporation into budding virions. Nevertheless, the precise localization of L and determination of the number of L incorporated into budding virions warrant future studies.

Interestingly, our FRAP data of viral condensates formed at late stages of EBOV infection show that late VFs exhibit a lower fluorescence recovery rate compared to early VFs (Fig. 6A), indicating that the viscosity of VFs progressively and dramatically increases during VF maturation. For EBOV, polymerase L, VP35 as well as RNA-binding by NP have been shown to influence the fluidity of reconstituted and infection induced viral condensates [8, 9]. Since the presence of condensed NCs is a main distinguishing feature between the early and late VFs, we hypothesized that NC condensation is responsible for the observed reduction in fluidity of late VFs. FRAP measurements of reconstituted early-like and late-like VFs demonstrated that VFs containing condensed NCs are less dynamic and more ordered (Fig. 6B-N). Hence, NC assembly and condensation is responsible for the reduced fluidity of EBOV VFs during maturation. Intriguingly, mumps virus VFs have been reported to undergo stress-induced coarsening [6] and measles VFs solidify upon ageing [40]. This highlights the possibility that NC assembly-driven reduced fluidity and/or solidification of VFs is a common feature among NNSVs and key to VF maturation marking the transition between replication (fluid phase) and genome packaging (gel-like phase). The importance of VF fluidity for viral replication is underscored by a recent study showing that replication of human respiratory syncytial virus is inhibited by a condensate hardening drug [41].

Taken together, our structural and FRAP data suggest that NC parallel bundles do not form solid crystalline arrays with long-range translational order, but may correspond to small paracrystalline domains separated by regions with lower order. As NCs undergo condensation, the rigidity of the NCs significantly increases. At the same time, free NP interfaces are depleted by assembly, which presumably decreases the VF volume anad hence increases the local concentration of condensed NCs. The combination of increased NC rigidity and concentration would be expected to drive parallel alignment of NCs. For moderate concentrations this would drive liquid-crystalline order; while above a threshold concentration and given sufficient relaxation time, the system would transition to crystalline order. One plausible scenario is that, as NC condensation proceeds the system initially acquires liquid crystalline order, and some regions of high local concentration form small crystalline domains. Whether the system exhibits liquid-crystalline and/or crystalline order needs to be addressed in future studies using mathematical modeling on capsid assembly. Our data suggest that the NC bundles become more ordered in time, and that they use this increase in physical order to disperse. As revealed by cryo-ET, VF dispersion of the late VFs leads to loss of condensate integrity and eventually allows interactions of the condensed NCs with cytoskeletal components. Upon condensation, we observed that NCs associate with short, filamentous structures with diameters similar to those of actin filaments (Fig. 4H-J, Movie S3). NCs stay associated with these filaments at all steps of virus budding, pointing towards actin-assisted transport and NC incorporation into budding virions. This data substantiates previous results indicating that long-distance transport of EBOV NCs is dependent on actin-polymerization [24, 42] and promoted by VP35 and VP24 [23], which localize to the appendages of condensed NCs (Fig. 4) [19, 21]. Hence it is plausible that the interaction of NC bundles with cytoskeletal components facilitates the separation of the bundles into individual NCs prior to virus budding at the plasma membrane. Since VP40 is also present at the plasma membrane and interacts with actin (Fig. S6) [29], it is likely that the actin network coordinates both the trafficking and incorporation of NCs into budding virions. Further studies are needed to determine the actin interaction site on the NC surface.

Taken together, our data allows us to propose a mechanistic model for EBOV replication and assembly (Fig. 8) which could be applied to other NNSV that form biomolecular condensates and show high NC assembly polymorphism [43, 44]. Our study reveals that VFs are governed by biophysical processes such as phase separation and increase in viscosity as NCs assemble, condense, and form bundles resembling liquid crystals. We demonstrate that assembly and alignment of NCs can lead to loss of integrity and dispersion of the condensate into the cytoplasm. This serves as a switch that allows the condensate material to interact with cellular components such as the cytoskeleton and can drive sudden changes in the cellular environment. In case of EBOV infection, the structural transition of the replication-competent loosely coiled NC into the condensed state drives alignment of NCs and a concomitant reduction in fluidity of the viral condensate. Eventually this leads to VF dispersion, loss of VF integrity, and accessibility of cytoskeletal components to the NCs. Finally, this allows for actin-dependent NC trafficking to the side of virus budding (Fig. 8). Overall, targeting viral condensates has a considerable potential for the development of drugs with broad-spectrum activity against emerging viral pathogens. We propose that condensate targeting drugs which would prevent dispersion or lead to premature dispersion and collapse of viral condensates may effectively antagonize RNA replication and expose viral RNA to recognition by the immune system.

**Fig. 8.**
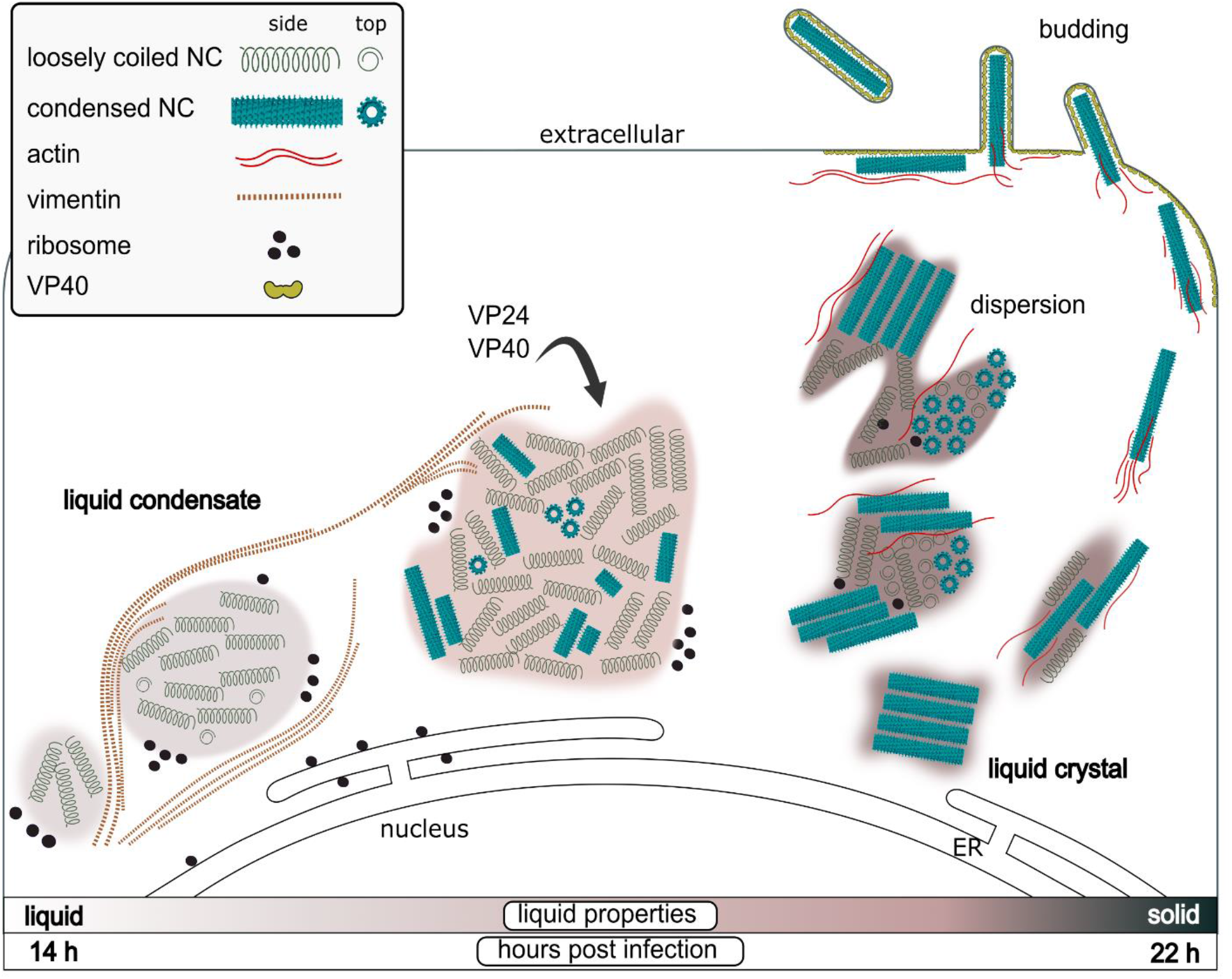
Proposed model for EBOV viral factory maturation. Early during infection, viral factories (VFs) form via liquid-liquid phase separation and harbor only loosely coiled NCs. VFs interact with the vimentin network which supports the formation and integrity of VFs. Upon recruitment of viral proteins VP24 and VP40 into the viral condensates, NCs undergo structural transition into condensed helical assemblies encapsidating the viral RNA genome. NC condensation drives NC bundle formation and a concomitant reduction in VF fluidity leading to VF dispersion late in infection. This second transition leads to loss of VF integrity rendering the NCs accessible to cytoskeletal components. Association of condensed NCs with actin drives their transport to the plasma membrane and incorporation into budding virions.

## Materials and Methods

### Cells

Huh7 (human hepatocarcinoma cells, kindly provided by Prof. Ralf Bartenschlager, Heidelberg University Hospital and Prof. Stephan Becker, Philipps-University Marburg) and HEK293T (Collection of Cell Lines in Veterinary Medicine CCLV-RIE1018) were maintained in Dulbecco’s Modified Eagle’s medium (DMEM, ThermoFisher Scientific) supplemented with 100 U/ml penicillin–streptomycin (ThermoFisher Scientific) and 10% (v/v) fetal bovine serum for maintenance (DMEM10) or 5% (DMEM5) for infection experiments at 37 °C and 5% CO_2_.

### Viruses and plasmids

Recombinant viruses were based on Ebola virus (EBOV) rec/COD/1976/Mayinga-rgEBOV (GenBank accession number <u>KF827427.1</u>, rgEBOV) [45]. The recombinant rgEBOV-VP30-GFP encoding a VP30 with GFP fused to its C-terminus, and rgEBOV-iluc2, carrying a codon-optimized firefly luciferase reporter have been described elsewhere [28, 33]. All work with infectious virus was performed under biosafety level (BSL-)4 conditions at the Friedrich-Loeffler-Institut (Federal Research Institute of Animal Health, Greifswald Insel-Riems, Greifswald, Germany) following approved standard operating procedures. Expression plasmids for the EBOV (Mayinga strain) proteins have been described elsewhere [28, 46].

### Infection of cells for structural analysis

For structural characterization of EBOV-infected cells, Huh7 cells were seeded on 200 mesh Quantifoil^TM^ SiO_2_ R1.2/20 or R1/4 EM grids placed on 3D-printed grid holders in a 96-well plate. 4-5 h after seeding of Huh7 cells at a seeding density of 7.5 x 10^3^ cells/well the samples were transferred to the BSL4 laboratory. Cells were infected with rgEBOV-WT, rgEBOV-VP30-eGFP at an MOI of 1 (based on the 50% tissue culture infectious dose [TCID50/ml]) in volumes of 100 µl. Cells were incubated for 14 h, 18 h, or 22 h before they were chemically fixed for 24 h with 4 % paraformaldehyde and 0.1 % glutaraldehyde in PHEM buffer (60 mM PIPES, 25 mM HEPES, 2 mM MgCl_2_, 10 mM EGTA, pH 6.9). 24 h after fixation the buffer was replaced with fresh fixation buffer and samples were fixed for another 24 h. After transfer of the samples out of the BSL4 containment, the grids were stored in PHEM buffer and plunge-frozen within three days.

### Transfection of cells for structural analysis

Holey silicon dioxide film coated grids (200 mesh Quantifoil^TM^ SiO_2_ R1.2/20) were placed into PDMS-coated dishes and were plasma-cleaned for 10 s in a Gatan Solarus 950 (Gatan). The dish was sterilized with 70% ethanol and washed twice with DMEM. After removal of the medium, cells were seeded on the grids at a seeding density of 1.8 × 10^5^ cells/dish. 4h post seeding, cells were transfected with plasmids in two different conditions: (1) 85 ng VP30-GFP, L, NP, VP35, VP24 and VP40 to facilitate formation of IBs containing condensed NCs (late-like IBs); (2) 125 ng VP30-GFP, L, NP, VP35 to induce the formation of IBs containing only loosely coiled NCs (early-like IBs). TransIT-LT1 transfection reagent (Mirus Bio LLC) was used according to manufacturer’s protocol. At 28 h post-transfection (p.t.) cells were vitrified by plunge-freezing as described below.

### Sample preparation for cryo-electron tomography

Chemically fixed EBOV-infected or transfected Huh7 cells on EM grids were vitrified by plunge-freezing into liquid ethane at −183 °C using a GP2 plunge-freezer (Leica) with a chamber temperature of 25 °C and 85% humidity. An additional 3 µl PHEM buffer (infected cells) or DMEM10 (transfected cells) were added to the grids before blotting them from the back with a Whatman Type 1 paper for 3 sec. Grids were clipped into FIB-AutoGrids^TM^ (ThermoFisher Scientific).

Cryo-focused ion beam (FIB) milling was performed using an Aquilos dual-beam cryo-focused ion beam-scanning electron microscope (cryo-FIB-SEM) (ThermoFisher Scientific) equipped with a cryo-stage cooled to −180 °C. Grid were loaded on a 45°-pretilt shuttle, cells were selected for milling and coated with organometallic platinum via the Gas Injection System and an injection time of 5 s. Cryo-lamellae were prepared by semi-automated milling in five (3+2) successive steps at a stage angle between 15 °C and 18 °C using a gallium-ion beam at an acceleration voltage of 30 keV and a modified version of the autolamella program available at https://github.com/DeMarcoLab/autolamella [47]. The first three steps were carried out automatically using the autolamella program, and the last two steps were performed manually with a nominal thickness of 150 nm. After polishing, lamellae were thinned at the back using a stage tilt of 21-22°.

### Cryo-ET image acquisition and data processing

Cryo-TEM montages and tilt series were collected on a Titan Krios Transmission Electron Microscope (TEM, ThermoFisher Scientific) operated at 300 keV and equipped with a BioQuantum® LS energy filter with a slit width of 20 eV and K3 direct electron detector (Gatan) using SerialEM [48]. Tilt series were acquired at 33,000× magnification with a pixel size of 2.671 Å/px at 2.5–4 µm defocus, with an electron dose of approximately 3 e^-^/Å^2^ per projection. A dose-symmetric acquisition scheme [37] was used with an effective tilt range of +68° to −52° in 3° increments.

Motion-correction was performed using MotionCor2 [49] and subsequent tomographic reconstruction was done using IMOD [50] or AreTomo[51]. Stacks of tomograms acquired on lamellae were aligned using patch tracking. Following 3D contrast transfer function (CTF) correction and dose-filtration implanted in IMOD, the tomographic reconstruction was performed by weighted back-projection with a simultaneous iterative reconstruction technique (SIRT)-like filter equivalent to 5 iterations. 3D segmentation and data analysis was performed using Dragonfly software, Version 2022.2.0.12227 (for Linux). Comet Technologies Canada Inc., Montreal, Canada; software available at https://www.theobjects.com/dragonfly.

### Subtomogram averaging

Subtomogram averaging of the EBOV NC was performed using the Dynamo software package 1.1.514 [52, 53]. Particles were picked using the crop along axis model with a subunit dz of 10 and dphi of 30 and subtomograms were extracted with a box size of 256 pixels. An initial template model was created by averaging of approximately 250 particles. To obtain an initial average, particles were aligned against the initial template using a Gauss filter smoothened cylindrical mask without imposing any symmetry. The initial average was then used as a template for the final averaging of 4,426 particles from 8 tomograms. The conical, azimuthal, translational search ranges and angular increments were gradually decreased within 5 iterations using parameters for refinement of the initial average (Supplementary Table S2). Adaptive filtering was used during the refinement. The attained resolution was estimated by comparing the two separately computed averages using FSC at 0.143 cut-off.

### Cryo-confocal microscopy of FIB lamellae and cryogenic on-lamella CLEM

Cells grown on EM grids were infected with rgEBOV-VP30-GFP and fixed 14 h, and 22 h p.i. and vitrified by plunge freezing. Cells were thinned by cryo-FIB milling and cryo-lamellae were imaged using a Zeiss LSM 900 Airyscan microscope equipped with a Linkam cryo-stage. To localize the cryo-lamellae, an overview image of the grid was acquired using a 5× air objective in widefield mode. Next, z-stacks with 0.3 µm spacing covering 3–5 µm were acquired in Airyscan mode SuperResolution with 488 and 405 laser lines at 2% and 640 at 0.02% using a Zeiss EC Epiplan-Neofluar 100× /0.75 DIC objective and a pixel size of 0.07 µm. In addition, reflection mode images were acquired for each z-plane. Z-stacks were 2D Airyscan processed using the Zen blue (ZEISS) build-in function and maximum intensity projections were generated in ImageJ/Fiji[54]. Subsequent correlation of the Airyscan images and low magnification TEM montages (lamellae maps) was performed in the eC-CLEM plugin [55] of Icy software using the lamellae shape and features such as lipid droplets visible in the reflection images guiding marks.

### Withaferin A treatment

To test the influence of Withaferin A (WFA) (abcam; ab120644) on VF formation of infected cells, Huh7 cells were seeded on 8-well chamber slides (Ibidi) (5 x 10^4^ cells/well) and 24 h later infected with rgEBOV-WT at an MOI of 3 (based on the TCID50/ml) in volumes of 100 µl or left untreated (mock) and incubated at 37 °C. 1 h p.i. the inoculum was removed, cells were washed with 200 µL PBS and fresh DMEM containing WFA at a final concentration of 6.25 µM was added to the cells or cells were left untreated (infected DMSO control and mock DMSO control). The cells were incubated for 14 h, 18 h or 22 h at 37 °C until they were chemically fixed for 24 h with 4 % paraformaldehyde and 0.1 % glutaraldehyde in DMEM5. Afterwards, cells were transferred out of the BSL4 containment and subjected to immunofluorescence analysis.

To test the influence of WFA on EBOV replication, HEK293T cells were seeded on 12-well plates (4 x 10^5^ cells/well) and 24 h later infected with rgEBOV-iluc2 at an MOI of 5 in volumes of 150 µl DMEM with or without WFA at finals concentrations of 1.5 and 6.25 µM or were left untreated (infected DMSO control and mock DMSO control) and incubated at 37 °C. 24 h later the cells were washed into the inoculum, which was then spun down for 5 min at 120 x g. Pellets were dissolved in 300 µl 1x Glo Lysis buffer (Promega) and incubated for 10 min at RT. Lysates were cleared by 3 min centrifugation at 10000 x g, and 50 µl cleared lysate was mixed with 50 µl Bright Glo reagent (Promega) for determination of viral RNA synthesis/protein expression or 50 µl CellTiter-Glo reagent (Promega) for determination of cell viability/ATP concentration. Luminescence was measured in white bottom plates using a TECAN Infinite F200 PRO microplate reader.

### Immunofluorescence analysis of infected and transfected cells

Fixed infected or transfected cells were washed three times with phosphate-buffered saline (PBS) before they were permeabilized in PBS containing 0.1% Triton X-100 for 10 min at room temperature. Subsequently, cells were washed with PBS and incubated for 1 h with blocking buffer (3% BSA in PBS supplemented with 0.1% Tween-20 (PBS-T). Cells were incubated with primary antibodies diluted in PBS-T with 1% BSA for 1 h at 4 °C on a rocker. After 3 washing steps with PBS-T for 5 min each, cells were incubated with secondary antibodies for 1 h at 4 °C on a rocker. After 3 washing steps with PBS-T for 5 min each, nuclei were fluorescently labeled with 1 mg/ml Hoechst 33342 (Sigma-Aldrich) in PBS for 5 min. Cells were washed 3 times with PBS for 5 min each, followed by a short washing step with deionized water. Coverslips were mounted on microscopy slides using 7 µl ProLong Glass Antifade Mountant (ThermoFisher Scientific, Invitrogen).

For samples shown in Fig. 1A polyclonal rabbit anti-Ebola virus NP (1:2000) (Gentaur) was used as primary antibody, and Alexa Fluor 488 goat anti-rabbit (1:1000) (Invitrogen) as secondary antibody.

For samples shown in Fig. 5F and S4D, polyclonal rabbit anti-Ebola virus NP (1:2000) (Gentaur), and anti-vimentin chicken polyclonal antibody (1:500) (PA1-10003, ThermoFisher Scientific) were used as the primary antibodies, Alexa Fluor 488 goat anti-rabbit (1:1000) (Invitrogen), and Alexa Fluor 647 goat anti-chicken IgG (H+L) (1:1000) were used as the secondary antibodies.

### Fluorescence data analysis and 3D segmentations

Airyscan processed fluorescence microscopy data was further analysed using Imaris software (Version 31 9.8.2, Oxford Instruments). Green or red fluorescent signal of IBs and vimentin respectively was segmented in three dimensions using the surface segmentation algorithm of Imaris. VFs of four cells from each time point (14, 18, 22 h p.i.) were segmented and the volume and sphericity were determined using the Imaris statistics function. The sphericity of each segmented IB, defined as the ratio of the surface area of a sphere with the same volume as the given VF to the surface area of the VF, was calculated using the Imaris statistics function.

### Fluorescence recovery after photobleaching (FRAP) analyses

#### FRAP analysis of reconstituted EBOV VFs

cells were seeded in 35 mm dishes (IBIDI) (3 x 10^5^ cells/dish), and 24 h later transfected with plasmids in two different conditions: (1) 85 ng VP30-GFP, L, NP, VP35, VP24 and VP40 to facilitate the formation of IBs containing condensed NCs (“late like IBs”); (2) 125 ng VP30-GFP, L, NP, VP35 to induce the formation of IBs containing only loosely coiled NCs. Fluorescence recovery was analysed 28 hours post transfection following Hoechst staining (1:1000) and medium change to FluoroBrite DMEM using a ZEISS laser scanning microscope (LSM900) equipped with an Airyscan detector and the ZEN Blue 3.1 software. Cells were kept in caged incubator maintaining 37 °C and 5% CO_2_. VP30-GFP and Hoechst fluorescence was excited with a 488 nm and 405 diode lasers at 0.6% laser power, respectively, and detected with a Plan-Apochromat 63× 1.4NA oil immersion objective (Zeiss) and with variable dichroic mirrors (410-480 nm for Hoechst and 500-550 nm for GFP detection). Photobleaching of whole IBs was performed using a 488 nm diode laser at 100%. One z-stack was recorded pre-bleach and post-bleach images were recorded in 10 second intervals for 25 minutes under the airyscan SR-4Y488 mode using a frame rate of 295.49 ms per frame and 488 nm and 405 diode lasers at 0.6% laser power. The z-stack range was set to 6 μM with 0.5 μM spacing. All images and stacks were Airyscan processed using the Zen blue (ZEISS) build-in function. The ImageJ/FIJI plugin MultiStagReg available at https://github.com/miura/MultiStackRegistration was used to align stacks of time series [54]. The mean fluorescence intensity of the first post-bleached VF spot was set to 0% and relative values for all frames were calculated. Average values were calculated for a total of 24 IBs of each of the conditions from three independent experiments.

#### FRAP analysis of VF in EBOV infected cells

FRAP analysis was done as previously described [8]. Briefly, Huh7 cells were seeded in 4-well chamber slides (IBIDI), and cells were infected with rgEBOV-VP30-GFP at an MOI of 1. One h p.i., medium was exchanged to DMEM with 5% FCS. Fluorescence recovery was analyzed 14, 18 and 22 h p.i. using a VisiScope Live Cell Imaging System with a 63x water immersion objective (Visitron Systems). Whole IBs were photobleached using a 405 nm laser (10 mW), with the FRAP time per pixel set to 100 ms. Images were taken in 10 second intervals for 5 minutes, and image analysis was performed using the VisiView 4.3.0.1 (Visitron Systems) software.

### High pressure freezing, freeze substitution and ultramicrotomy

Sapphire discs with a diameter of 3.0 mm and 50 μm thickness (Wohlwend GmbH) were cleaned with ethanol (100%) and coated with a 15 nm carbon layer using a sputter coater (EM ACE600, Leica). To distinguish between the two sides of the disc, the letter “F” was engraved on the carbon side. Afterwards, the discs were incubated at 120 °C overnight. For transfection experiments, polydimethylsiloxane (PDMS) coated 35 mm dishes and the carbon-coated sapphire discs were plasma-cleaned for 10 seconds (s) in a Solarus 950 (Gatan). The discs were placed inside the dishes with the “F” facing up, sterilized in 70% ethanol and washed twice with fresh DMEM medium. After removal of DMEM, Huh7 seeded on the sapphire discs at a seeding density of 0.22 x 10^6^ cells/dish, followed by transfection 6 hours post seeding using different EBOV plasmids and TransIT-LT1 transfection reagent (Mirus Bio LLC) according to manufacturer’s protocol. 48 h post transfection, cells were processed by high pressure freezing (HPF) and freeze substitution (FS) as described below.

For infection experiments, carbon-coated sapphire discs were placed on 3D-printed grid holders in a 96-well plate, sterilized in 70% ethanol and washed twice with DMEM. 7.5 x 10^3^ Huh7 cells per well were seeded and the plates were transferred to the BSL4 laboratory after 4-5 h. 24 h post seeding, the cells were infected with rgEBOV-WT at an MOI of 1. At 14, 18, or 22 h p.i., the infected cells were washed with PBS and fixed with 4% formaldehyde and 0.1% glutaraldehyde in PHEM buffer (60 mM PIPES, 25 mM HEPES, 10 mM EGTA, 2 mM MgCl_2_) for 24 hours and stored in buffer until further processing for a maximum of three days.

Sapphire discs with either transfected or infected cells were assembled between two 1-hexadecene-treated specimen carriers (Type A and B, Wohlwend GmbH) with cells facing the 100 µm deep cavity of carrier Type A. Vitrification of cells was done by high-pressure freezing at a minimum of 2,200 bar maintained for 370 ms and a cooling rate of 20,000 K/s using a Leica EM ICE. After transfer of the sapphire discs from liquid nitrogen to a freeze-substitution (FS) solution (0.1% uranyl acetate in anhydrous acetone) the samples were cooled to −90 °C and processed in an automated FS system (EM AFS2, Leica). After a washing step in acetone, samples were infiltrated with Lowicryl HM20 (Polysciences Inc) and polymerized using UV light. For the detailed FS protocol see Table S2. Sectioning of the lowicryl-embedded samples was done using diamond knifes (DiATOME) and an ultramicrotome (UC7, Leica). Sections with 100 nm nominal thickness were collected on 2 × 1 mm copper slot grids (Gilder) coated with pioloform support film.

### Immunogold labeling of resin sections

Immunogold labeling for detection of EBOV NP within the VFs and VP40 at the budding sites was performed using polyclonal rabbit anti-EBOV NP antibody (Gentaur, 0301-012) or polyclonal rabbit anti-EBOV VP40 (IBT Bioservice, 0301-010) and 10 nm gold-conjugated protein A (Aurion, 110.111). To confirm the presence of vimentin, sections were incubated with anti-vimentin mouse antibody (Santa Cruz Biotechnology, sc-6260), followed by staining with a bridging rabbit anti-mouse antibody (Life Technologies, A27022) and 10 nm gold-conjugated protein A (Aurion, 110.111).

### Electron microscopy of resin sections

Grids were imaged with a Talos L120C TEM operated at 120 keV and equipped with a LaB_6_ filament and a Ceta-M camera with a 4k × 4k CMOS (Thermo Fisher Scientific). Electron micrographs were recorded at magnifications of 5,300 ×, 13,500 ×, and 45,000 × (corresponding pixel sizes at the specimen level: 26.44 Å, 13.35 Å and 3.28 Å respectively) using target counts between 2500 and 3000. Mapping and data acquisition was performed in SerialEM [48].

### Statistical analyses

Statistical analyses were performed in Graphpad Prism 9.5.0. The details of the quantification and all statistical analyses are included in figure captions or the relevant sections of material and methods section.

## Data availability

Electron tomography data have been deposited to the Electron Microscopy Data Bank under accession codes EMD-18675 (tomogram of Fig. 3K), EMD-18676 (tomogram of Fig. 3F), EMD-18678 (tomogram Fig. 5A), EMD-18679 (tomogram of Fig. 6C), EMD-18686 (tomogram of Fig 6I), EMD-18690 (tomogram of Fig. 2C), EMD-18695 (tomogram of Fig. 2I), EMD18696 (tomogram of Fig 3A), EMD-18697 (subtomogram average of the nucleocapsid) and will be available upon publication. Additional data and material related to this publication may be obtained upon request.

## Acknowledgments

We thank Dr. Vibor Laketa and Dr. Sylvia Olberg for instructions on FRAP. We thank the Infectious Diseases Imaging Platform (IDIP) at the Center for Integrative Infectious Disease Research Heidelberg, the cryo-EM network at the Heidelberg University (HDcryoNET) and Heidelberg University Electron Microscopy Core Facility for support and assistance. The authors gratefully acknowledge the data storage service SDS@hd supported by the Ministry of Science, Research, and the Arts Baden-Württemberg (MWK), the German Research Foundation (DFG) through grant INST 35/1314-1 FUGG and INST 35/1503-1 FUGG. We thank Dr. Simone Mattei and Dr. Zhengyi Yang (EMBL Heidelberg) for training on cryo-confocal microscope Zeiss Airyscan. We are grateful for the discussion and critical reading of the manuscript to Prof. Ulrich Schwarz and Prof. Michael Hagan.

## Author contributions

Conceptualization: MV, TH, PC; Methodology: MV, BSB, KF, JM, SLW, MWM, TH, PC; Investigation: MV, BSB, KF, JM, LW, TH; Visualization: MV; Funding acquisition: MV, PC; Project administration: PC; Supervision: TH, PC, MV; Writing – original draft: MV, PC; Writing – review & editing: MV, BSB, KF, SLW, MWM, LW, TH, PC

