## Supplementary material for "Nucleocapsid condensation drives Ebola viral factory maturation and dispersion"

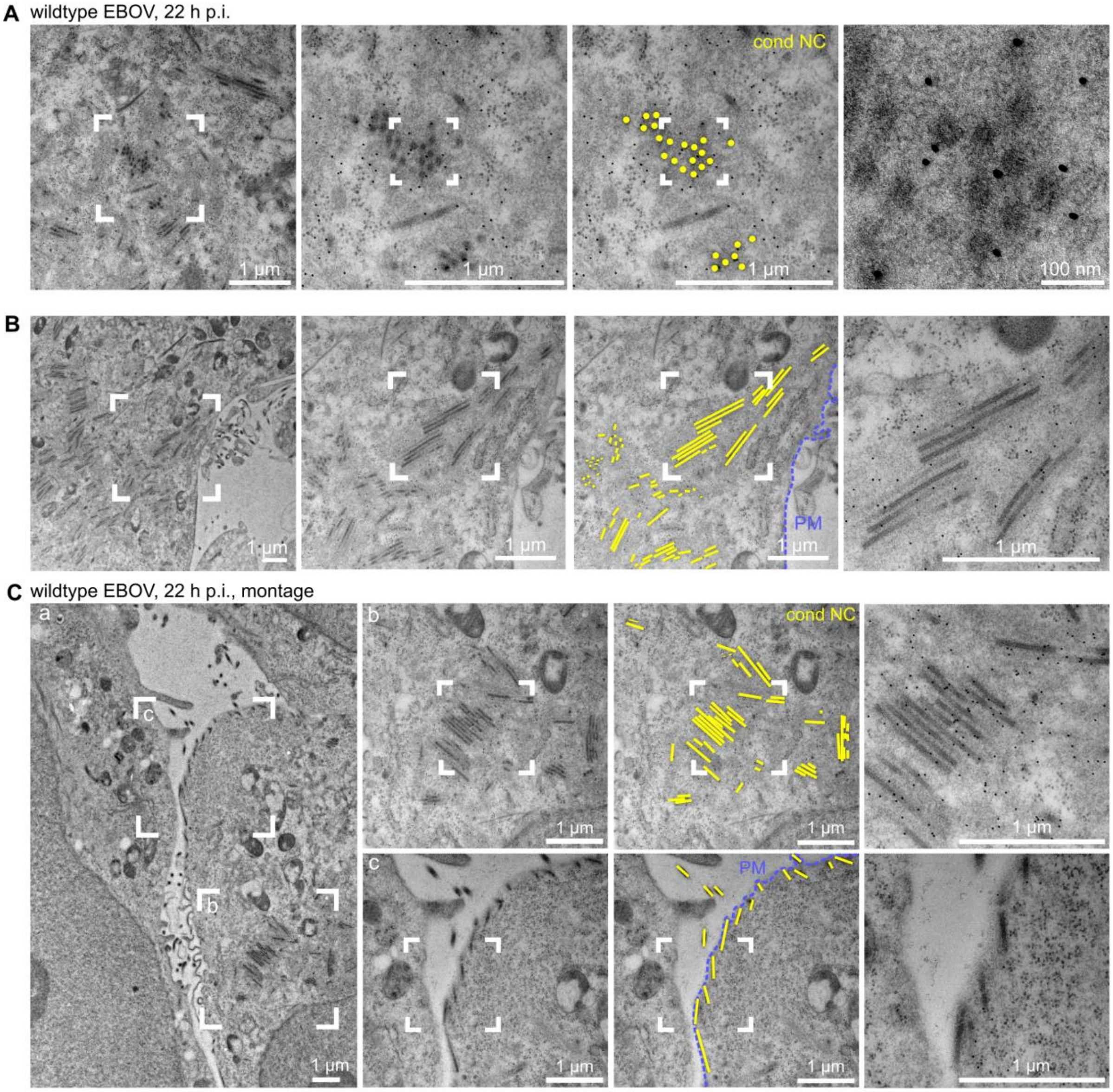


### Figure S1

### Ebola virus nucleocapsids form large bundles that separate near the plasma membrane

**(A-C)** Thin-section transmission electron microscopy analysis of EBOV viral factories (VFs) at 22 h p.i. with wildtype EBOV. VFs were identified by immuno-labelling of EBOV NP. Sections: 100 nm. Top **(A)** and side view **(B)** of condensed nucleocapsids (cond NCs, yellow) organized into locally ordered bundles near the plasma membrane (PM, blue). Montage image and close-up views showing Individual condensed NCs aligning at the PM **(C)**.

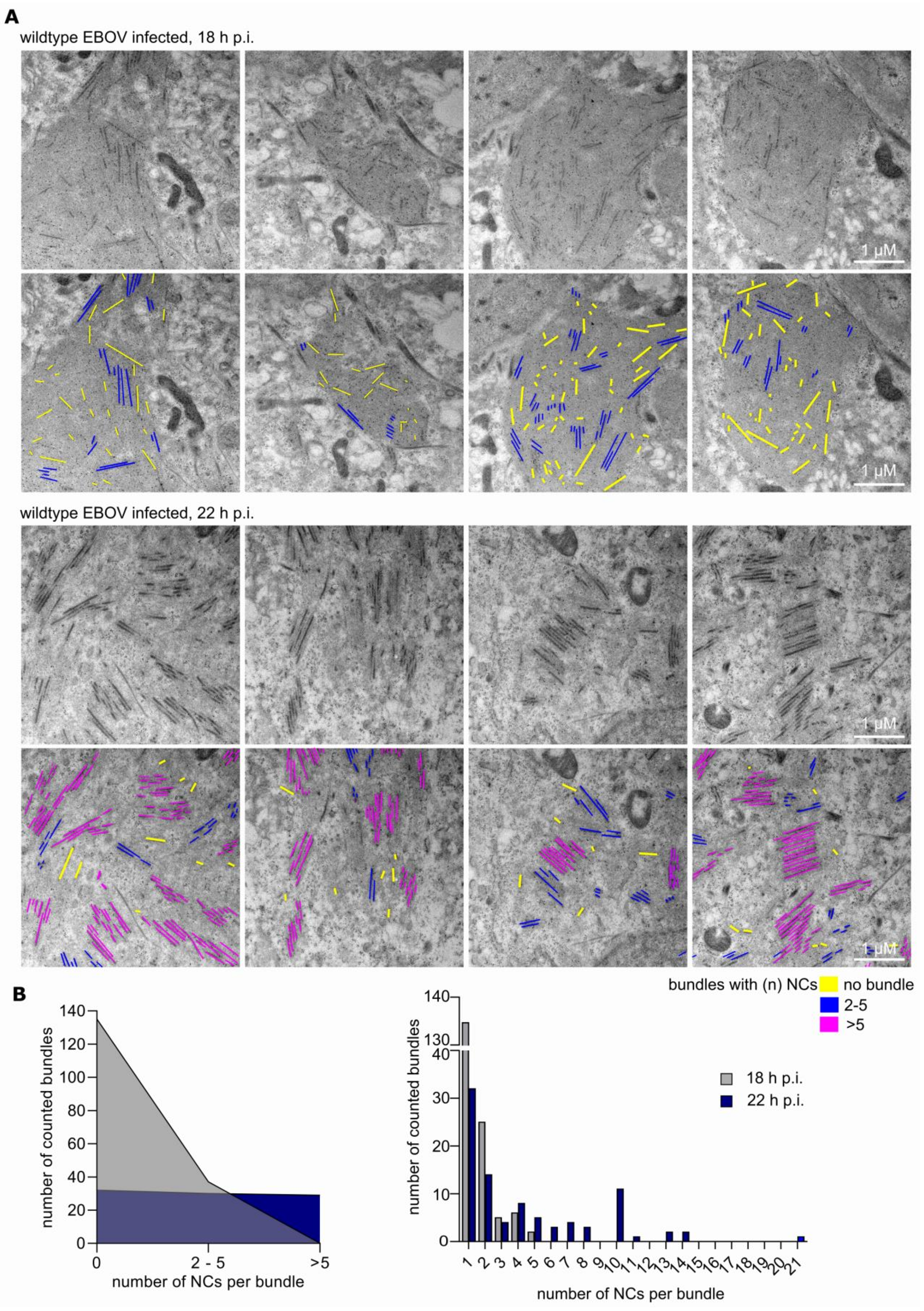
A
wildtype EBOV infected, 18 h p.i.
1 µM
wildtype EBOV infected, 22 h p.i.
B
bundles with (n) NCs
no bundle
2-5
>5
number of counted bundles
number of NCs per bundle
0
2 - 5
>5
18 h p.i.
22 h p.i.

### Figure S2

#### Ebola viral factory dispersion coincides with formation of large nucleocapsid bundles

**(A)** Thin-section transmission electron microscopy of EBOV VFs at 18 h p.i. (upper panel) and 22 h p.i. (lower panel) with wildtype EBOV. Representative images VFs from four 100 nm sections are shown. Individual condensed nucleocapsids (NCs) are highlighted in yellow, bundles of 2-5 NCs in blue and large bundles of more than 5 NCs per bundle in pink.

**(B)** Quantification of NC bundles counted per indicated time point. Left panel shows the total number of NC bundles with 2-5 or >5 NCs per bundles counted in the four images shown in **(A)**. Right panel shows the total number of NC bundles with 1-21 NCs per bundle.

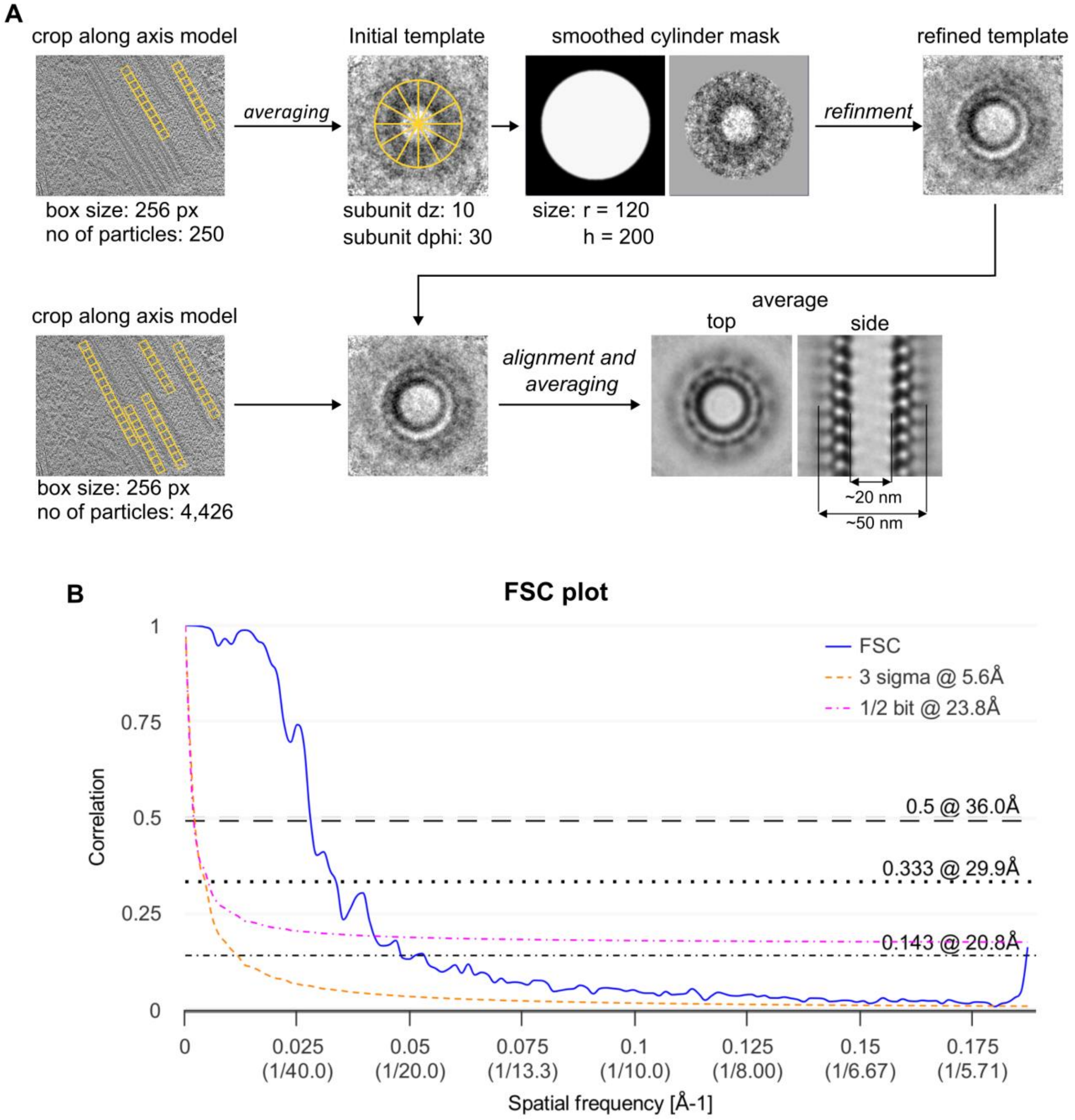


### Figure S3

### Workflow for subtomogram averaging and Fourier Shell Correlation plot

**(A)** Subtomogram averaging of the intracellular EBOV NC was performed using the Dynamo software package 1.1.514. Particles were picked using the crop along axis model with a subunit dz of 10 and dphi of 30 and subtomograms were extracted with a box size of 256 pixels. An initial template model was created by averaging of approximately 250 particles. To obtain an initial average, particles were aligned against the initial template using a Gauss filter smoothened cylindrical mask without imposing any symmetry. The initial average was then used as a template for the final averaging of 4,426 particles from 8 tomograms.

**(B)** Resolution estimation by Fourier Shell Correlation (FSC) of two independently calculated averages obtained by multiple alignment and averaging rounds using adaptive filtering in Dynamo. FSC was calculated using the Fourier Shell Correlation Server at the Electron Microscopy Data Bank (https://www.ebi.ac.uk/emdb/validation/fsc/results/) webpage using a smooth cylindrical mask, degree of symmetry 1 and filling degree 0.66.

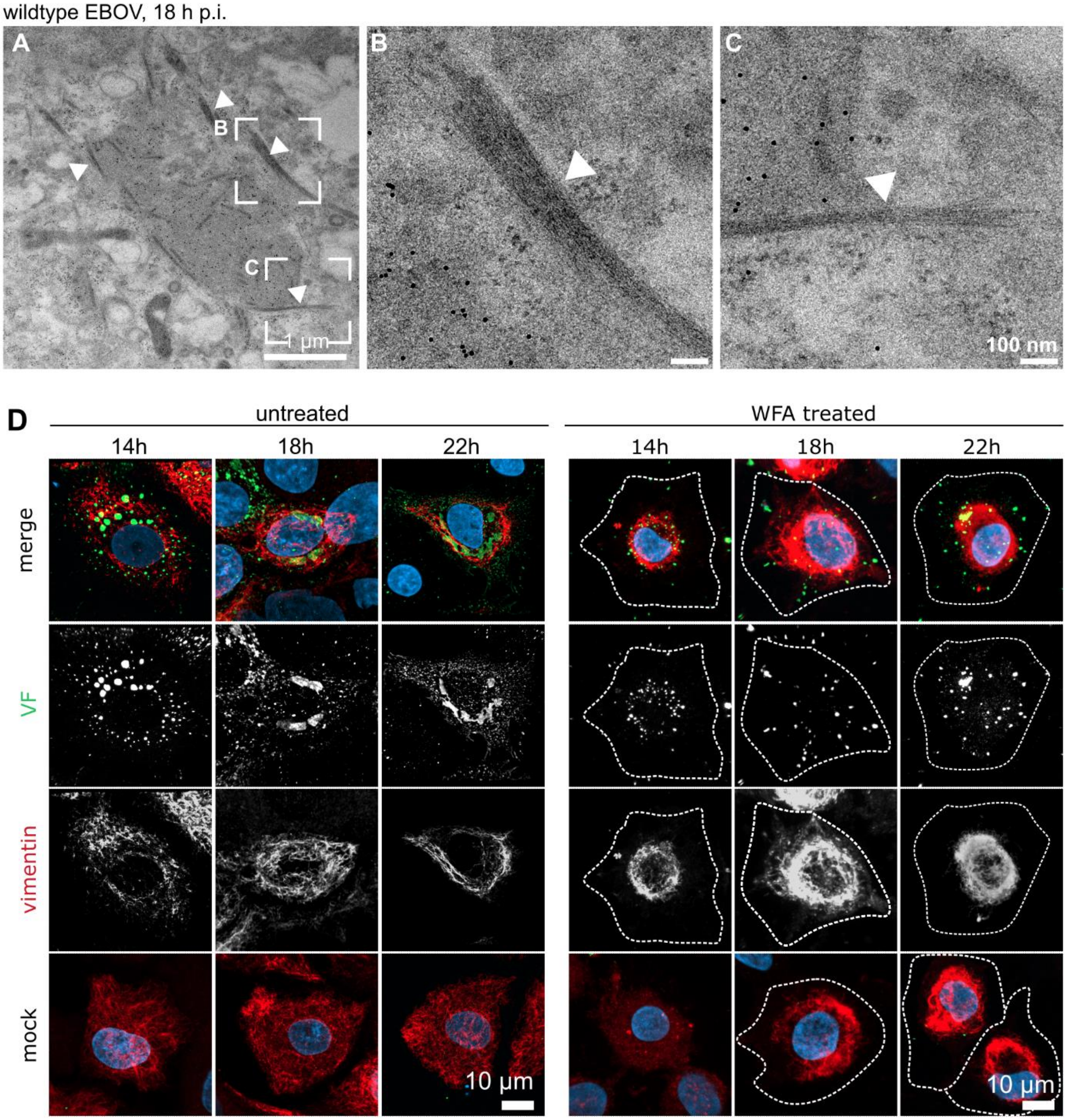


### Figure S4

### EBOV VFs are connected to large networks of vimentin intermediate filaments

**(A-C**) Thin-section transmission electron microscopy analysis of EBOV viral factories (VFs) at 18 h p.i. identified by immuno-labelling of EBOV NP. Arrow heads indicate intermediate filaments (IFs) associated with the VFs **(A)**. Close-up views of IFs. Arrow heads indicate IFs **(B,C)**.

**(D)** Confocal microscopy analysis of wildtype EBOV infected cells fixed at 14, 18 and 22 h p.i. Cells were immuno-stained against the EBOV nucleoprotein forming the viral factories (VFs) (green) and against vimentin intermediate filaments (red). Nuclei were stained with Hoechst (blue). Shown are threedimensional segmentations of the VF and vimentin (upper panel) and respective enlarged views (lower panels). Right panels show 3D segmentations of cells treated with withaferin A (WFA) at a concentration of 6.25 µM. Dashed line indicates the plasma membrane.

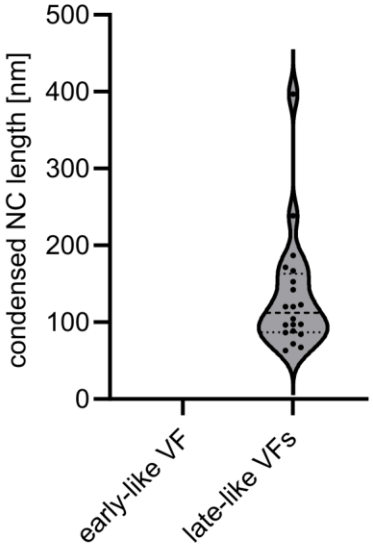


**Figure S5**

**Nucleocapsids in reconstituted EBOV VFs vary in length**

Length measurements of condensed NCs in cryo-lamellae prepared from cells 28 h after transfection with expression plasmids encoding EBOV NP, VP35, VP30-GFP and L (early-like VF) or additionally VP24 and VP40 (late-like VF). Violin plot shows the length distribution of condensed NCs in early-like (not detected) and late-like VFs (134 ± 75). n = 20 NCs measured in 5 tomograms acquired on three cryo-lamellae.

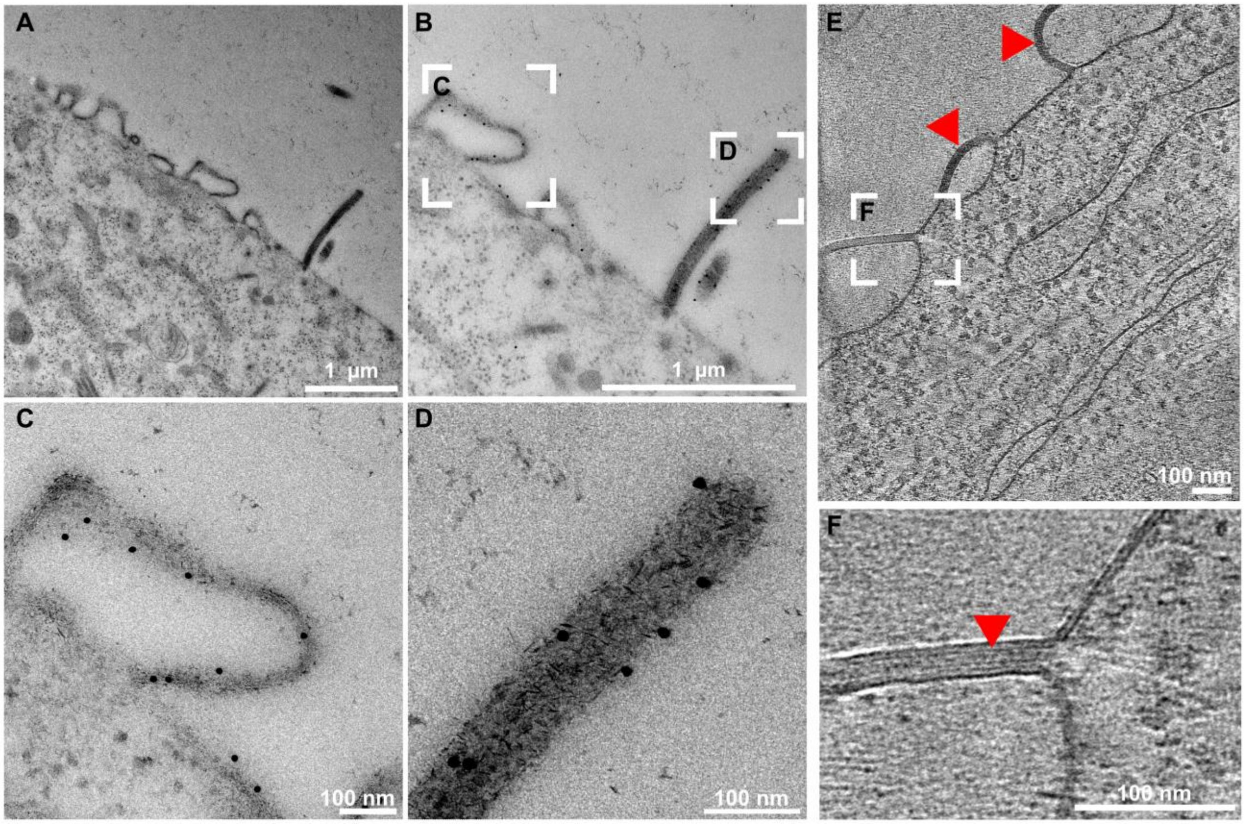


**Figure S6**

**EBOV VP40 localizes to viral budding sites**

**(A-D**) Thin-section transmission electron microscopy analysis of wildtype EBOV infected cells at 22 h p.i. EBOV VP40 was identified by immunogold labelling. Zoom-in into VP40 induced plasma membrane ruffles **(C)** and a budding virion **(D)**.

**(E, F)** *In situ* cryo-ET of wildtype EBOV infected cells at 22 h p.i. Tomogaphic slice showing plasma membrane ruffles containing VP40 **(E)** and zoom-in view showing the VP40 layer **(F)** with red arrows indicating VP40-containing membrane ruffles.

**Movie S1**

Movie S1 shows a tomogram of an EBOV VF that formed in a cell at 14 h p.i. with wildtype EBOV revealing that early VFs are mainly composed of loosely coiled helical NCs and exclude cellular material. Scale bar: 100 nm. Movie S1 is related to Figure 3A.

**Movie S2**

Movie S2 shows a tomogram of an EBOV VF that formed in a cell at 22 h p.i. with wildtype EBOV. It shows that late VFs are composed of both loosely coiled helical NCs and condensed NCs. The condensed NCs seen from top view organize in small hexagonal arrays forming parallel bundles. Scale bar: 100 nm. Movie S2 is related to Figure 3K.

**Movie S3**

Movie S3 shows a condensed EBOV NC in the cytoplasm outside of a VF and close to the plasma membrane. The NC is associated with thin filaments corresponding to actin. The tomogram was acquired on a lamella prepared from cells infected with wildtype EBOV and fixed at 22 h p.i. Scale bar: 100 nm. Movie S3 is related to Figure 4H.

**Movie S4**

Movie S4 shows an EBOV VF in a wildtype EBOV infected cell at 14 h p.i. revealing that VFs are associated with vimentin intermediate filaments that invade the space of the VFs. Scale bar: 100 nm. Movie S4 is related to Figure 5A.

**Movie S5**

Movie S5 shows a reconstituted EBOV VF formed in cells at 28 h post transfection with expression plasmids encoding EBOV NP, L, VP35 and VP30-GFP. The tomogram reveals that in the absence of VP24 and VP40 reconstituted VFs are mainly composed of loosely coiled NCs resembling early-like VFs. Movie S3 is related to Figure 6C.

**Movie S6**

Movie S6 shows a reconstituted EBOV VF formed at 28 h post transfection of cells with expression plasmids encoding EBOV NP, L, VP35, VP30-GFP, VP24 and VP40. The tomogram shows that in presence of VP24 and VP40 reconstituted VFs contain both loosely coiled NCs and condensed NCs and thus resemble late-like VFs. Movie S3 is related to Figure 6I.

**Table S1: Numerical parameters for subtomogram averaging in Dynamo.**

| Parameters | Round 1 | Round 2 | Round 3 | Round 4 | Round 5 |
|---|---|---|---|---|---|
| **iterations** | 1 | 1 | 1 | 1 | 1 |
| **references** | 1 | 1 | 1 | 1 | 1 |
| **cone aperture [°]** | 360 | 24 | 12 | 6 | 6 |
| **cone Sampling [°]** | 60 | 8 | 4 | 2 | 2 |
| **azymuth rotation range [°]** | 60 | 30 | 12 | 6 | 3 |
| **azymuth rotation sampling [°]** | 20 | 10 | 4 | 2 | 1 |
| **refine** | 5 | 5 | 5 | 5 | 5 |
| **refine factor** | 2 | 2 | 2 | 2 | 2 |
| **high pass filter [pixel]** | 2 | 2 | 2 | 2 | 2 |
| **low pass filter [pixel]** | 32 | 64 | 64 | 80 | 128 |
| **symmetry** | C1 | C1 | C1 | C1 | C1 |
| **particle dimensions** | 64 | 128 | 128 | 128 | 256 |
| **shift limits in X, Y and Z [pixel]** | 12, 12, 12 | 8, 8, 8 | 4, 4, 4 | 2, 2, 2 | 1, 1, 1 |
| **shift limiting way** | 2 | 2 | 2 | 2 | 2 |
| **separation in tomogram** | 1 | 1 | 1 | 1 | 1 |

**Table S2: Automated freeze substitution program.**

| step | T start | T end | Slope | Time | Reagent | Transfer | Agitation | UV |
|---|---|---|---|---|---|---|---|---|
| 1 | -90°C | -90°C | 0 | 12 h | 0.1% UA | stay | off | off |
| 2 | -90°C | -45°C | 7.5 °C/h | 6 h | 0.1% UA | stay | off | off |
| 3 | -45°C | -45°C | 0 | 25 min | acetone | exch/fill | off | off |
| 4 | -45°C | -45°C | 0 | 25 min | acetone | exch/fill | off | off |
| 5 | -45°C | -45°C | 0 | 25 min | acetone | exch/fill | off | off |
| 6 | -45°C | -45°C | 0 | 3 h | 25% Lowicryl | mix | on | off |
| 7 | -45°C | -35°C | 3.3 °C/h | 3 h | 50% Lowicryl | mix | on | off |
| 8 | -35°C | -25°C | 3.3 °C/h | 3 h | 75% Lowicryl | mix | on | off |
| 9 | -25°C | -25°C | 0 | 4 h | 100% Lowicryl | exch/fill | off | off |
| 10 | -25°C | -25°C | 0 | 6 h | 100% Lowicryl | exch/fill | off | off |
| 11 | -25°C | -25°C | 0 | 12 h | 100% Lowicryl | exch/fill | off | off |
| 12 | -25°C | -25°C | 0 | 24 h | 100% Lowicryl | stay | off | on |
| 13 | -25°C | 20°C | 3.7 °C/h | 12 h | 100% Lowicryl | stay | off | on |
| 14 | 20°C | 20°C | 0 | 12 h | 100% Lowicryl | stay | off | on |

UA: uranyl acetate; exch: exchange
